# Mapping single-cell responses to population-level dynamics during antibiotic treatment

**DOI:** 10.1101/2022.11.18.517151

**Authors:** Kyeri Kim, Teng Wang, Helena R. Ma, Emrah Şimşek, Boyan Li, Virgile Andreani, Lingchong You

**Affiliations:** Department of Biomedical Engineering, Duke University, USA; Center for Quantitative Biodesign, Duke University, USA; Integrated Science Program, Yuanpei College, Peking University, China; Biomedical Engineering Department, Boston University, USA; Biological Design Center, Boston University, USA; Center for Genomic and Computational Biology, Duke University, USA; Department of Molecular Genetics and Microbiology, Duke University School of Medicine, USA

**Author notes:** Correspondence and requests for materials should be addressed to Lingchong You.

## Abstract

Treatment of sensitive bacteria with beta-lactam antibiotics often leads to two salient population-level features: a transient increase in total population biomass before a subsequent decline, and a linear correlation between growth and killing rates. However, it remains unclear how these population-level responses emerge from collective single-cell responses. During beta-lactam treatment, it is well recognized that individual cells often exhibit varying degrees of filamentation before lysis. We show that the probability of cell lysis increases with the extent of filamentation and that this dependence is characterized by unique parameters that are specific to bacterial strain, antibiotic dose, and growth condition. Modeling demonstrates how the single-cell lysis probabilities can give rise to population-level biomass dynamics, which were experimentally validated. This mapping provides insights into how the population biomass time-kill curve emerges from single cells and allows the representation of both single-and population-level responses with universal parameters.

## Introduction

Beta-lactam antibiotics are widely used to treat bacterial infections ^1^. Studies have shown that beta-lactams exhibit both time- and antibiotic concentration-dependent killing ^2^. More specifically, two salient dynamic features of time-dependent killing were demonstrated in time-kill curves at a lethal antibiotic concentration. First, the total population biomass will first increase before collapsing, leading to an apparent time-delay in the effect of the antibiotic ^3, 4^. Second, the maximum lysis rate is linearly correlated to the growth rate of the population before antibiotic treatment ^5, 6^. Quantitative measurements of these dynamics have been shown to be important for designing effective antibiotic treatment protocols ^6, 7^. However, it is unclear how these population-level features could emerge from the collective responses of single bacterial cells undergoing antibiotic treatment ^2^.

At single-cell level, exposure to beta-lactam antibiotics often results in morphological changes in rod-shaped bacteria ^8-16^. Beta-lactams block cell wall synthesis by binding to penicillin-binding proteins (PBPs) and inhibiting peptidoglycan cross-linking ^17-19^, while the cells continue to accumulate biomass ^20, 21^. As a result, individual cells elongate exponentially without cell division and become filamented ^15^. Due to the loss of cell-wall integrity, filamentation is eventually followed by rapid cell lysis, consisting of cell membrane bulging through the cell wall pore and subsequent bursting of the cell membrane ^2, 22-27^. The time for each cell to lyse has been shown to vary under prolonged antibiotic exposure ^28^. Additionally, beta-lactam-induced filamentation can be reversible: if the antibiotic is removed before lysis occurs, filamented cells can divide into multiple cells ^10, 24, 29, 30^, with the resulting cell number being roughly proportional to the filament length ^31^. Therefore, bacterial filamentation affects population recovery ^12^. These observations underscore the importance in quantifying key parameters of cellular lysis to couple with biomass dynamics of the population.

Despite the evident connection between single-cell filamentation and population dynamics, the lack of quantitative analysis prevents a clear understanding of how the population dynamics emerge from the collective elongation and lysis of the single cells. Here, we found that the lysis probability increases with the extent of filamentation by measuring single-cell filamentation and lysis dynamics using time-lapse microscopy. We found that key parameters of the lysis probability are unique to bacterial strain and antibiotic dose. We further show how the single-cell elongation and lysis parameters can explain different, experimentally measured population dynamics resulting from beta-lactam treatment (**Figure 1a**). Our results provide new insights into the mechanism of beta-lactam-induced cell lysis and a better understanding of the coupling between single-cell- and population-level dynamics.

**Figure 1.**
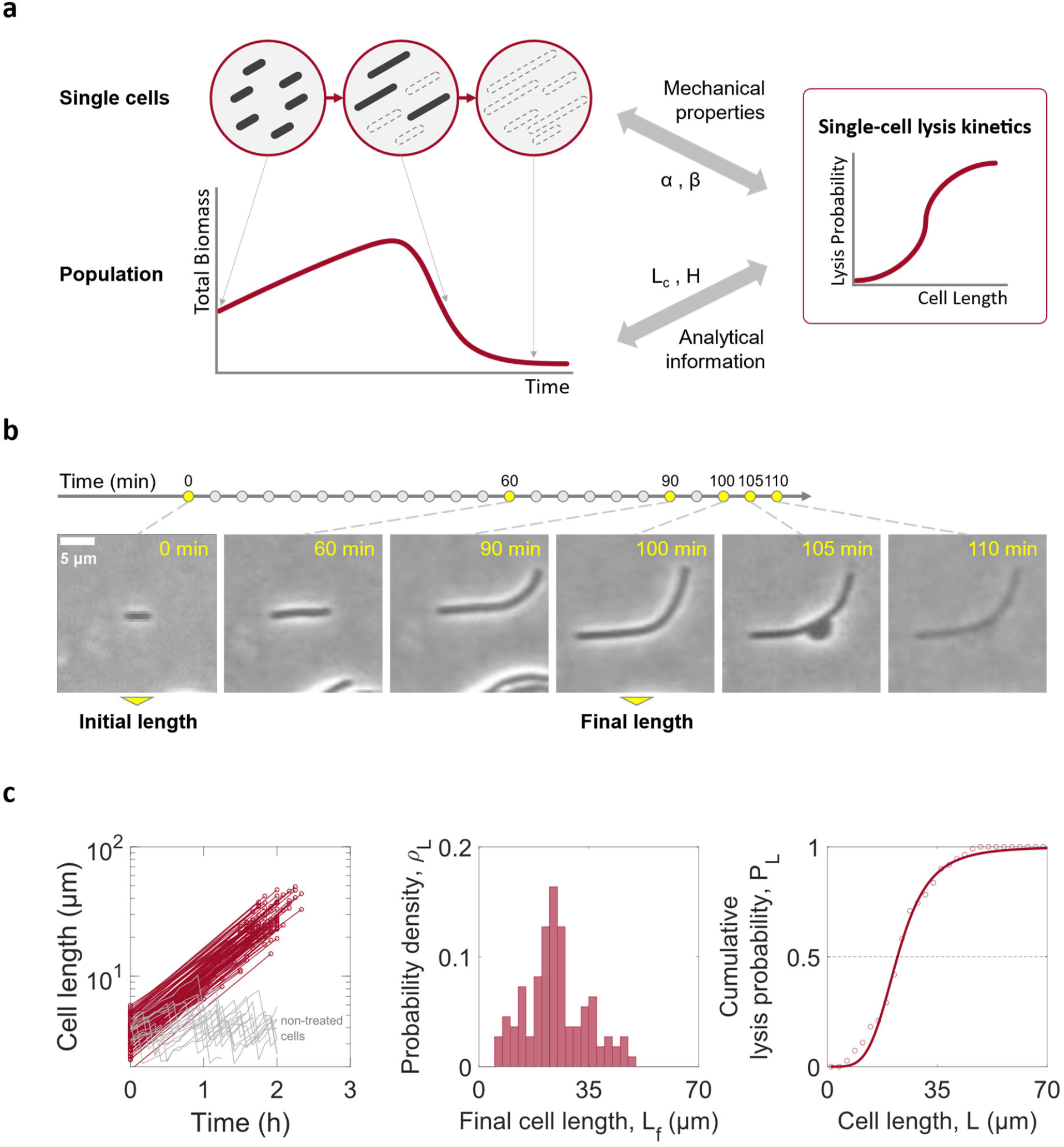
Mapping single-cell kinetics to population dynamics during beta-lactam treatment. **a. A schematic of single-cell versus population-level responses**. Bacteria elongate in response to beta-lactam antibiotics. During prolonged exposure, lysis occurs when both cell wall and membrane integrity are broken. At the population level, the total biomass, which is a sum of survivors’ biomass, experiences a transient increase before a decline due to single-cell filamentation and lysis. **b. Single-cell lengths were measured from time-lapse microscopy images upon beta-lactam antibiotic treatment**. Initial cell length was measured in the first time frame. Elongation ends with cell length shrinkage due to bulging. The longest length of each cell across the time frames was used as the final length. **c. Single-cell lysis probabilities were plotted with the extent of filamentation**. Red circle markers connected with lines show the measured initial and final lengths of 110 carbenicillin-treated individual cells over time, while gray lines show the length of 15 non-treated cells with division (left panel, log scaled in length). Probability density distribution (*ρ*_*L*_) over final cell lengths was collected by normalizing the probability distribution (middle panel). Dot plotted cumulative lysis probability distribution (*P*_*L*_) of the *ρ*_*L*_ is shown, and was fitted to the log-logistic distribution in cell length (right panel).

## Results

### The probability of antibiotic-induced lysis depends on filamentation length

We tracked beta-lactam-mediated filamentation of single *E. coli* MG1655 cells using time-lapse microscopy (**Figure 1b**). Cells in exponential growth phase were loaded onto a thin agarose gel containing growth medium and beta-lactam antibiotics. We tracked the cells and collected initial and final lengths over time (**Figure 1c**). We defined the final length as the longest length of a cell during observation in a given time interval for two reasons: 1) cells elongated only in the long-axis direction and shrunk when the cell wall burst ^23^, and 2) once bursting was observed, lysis took place in a short time ^22-24^. For example, we observed that filamentation of a cell was observed for 100 minutes, followed by bulging and lysis in the next 10 minutes, a period 10 times shorter than the duration of filamentation (**Figure 1b**).

Our measurements showed that lysis probability depends on the extent of elongation. To find single-cell lysis kinetics in cell length during elongation, we first measured the lysis probability density (*ρ*_*L*_) from the final lengths, which represents the fraction of cells that was lysed at each final length. We observed that *ρ*_*L*_ is small for both short cells and long cells and peaks at intermediate cell lengths. A small *ρ*_*L*_ at a small *L* reflects a small lysis probability for a short cell. A small *ρ*_*L*_ at a large *L*, however, does not imply a small lysis probability for a long cell, but rather the rarity of such cells due to lysis before reaching a long length. The shape of the dependence of *ρ*_*L*_ on *L* suggests the extent of elongation is predictive of the likelihood of lysis.

We next constructed the cumulative lysis probability (*P*_*L*_) from *ρ*_*L*_, i.e., 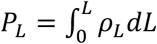, which is the probability for a cell to lyse before reaching length *L* (**Figure 1c**). We empirically chose to fit the sigmoidal dependence of *P*_*L*_ on *L* using a Hill equation (corresponding to a log-logistic distribution):

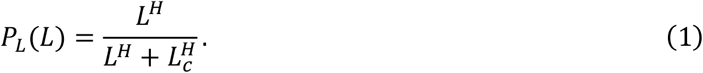

We term *L*_*c*_ the critical length; 50% of cells would lyse before reaching this length and *ρ*_*L*_ peaks at *L*_*c*_. The Hill coefficient, *H*, represents the steepness of *P*_*L*_. This fitted equation is phenomenological, the sigmoidal curve can be fit to other equations. However, as we shall see, the choice of the Hill equation provides a simple analytical interpretation of the population dynamics during antibiotic treatment. The dependence of lysis probability on extent of elongation is not unique to cells treated with a beta-lactam: it also applies to *E. coli* cells treated with another cell-wall synthesis inhibitor, D-cycloserine, according to our analysis of the raw data provided in a study ^19^ (**Figure S1**).

### The critical length depends on the antibiotic dose

The lysis probability exhibited the same dependence on filamentation length in different antibiotic and growth conditions. We exposed bacteria to three different carbenicillin concentrations (20, 50, and 100 μg/ml) and incubated at two different temperatures (27 and 37 °C) (**Figure 2a** and **Figure S2**). Temperature changed elongation rate but did not affect *L*_*c*_. However, *L*_*c*_ decreased with increasing carbenicillin concentration (**Figure 2b**). This inverse correlation confirms that cells can better tolerate lower-dose antibiotics and elongate further. Moreover, the correlation between initial and final lengths was weak under all conditions; that is, the elongation capacity was not defined by initial length during antibiotic exposure (**Figure S2a**). *H* did not show a dependence on antibiotic dose or temperature changes.

**Figure 2.**
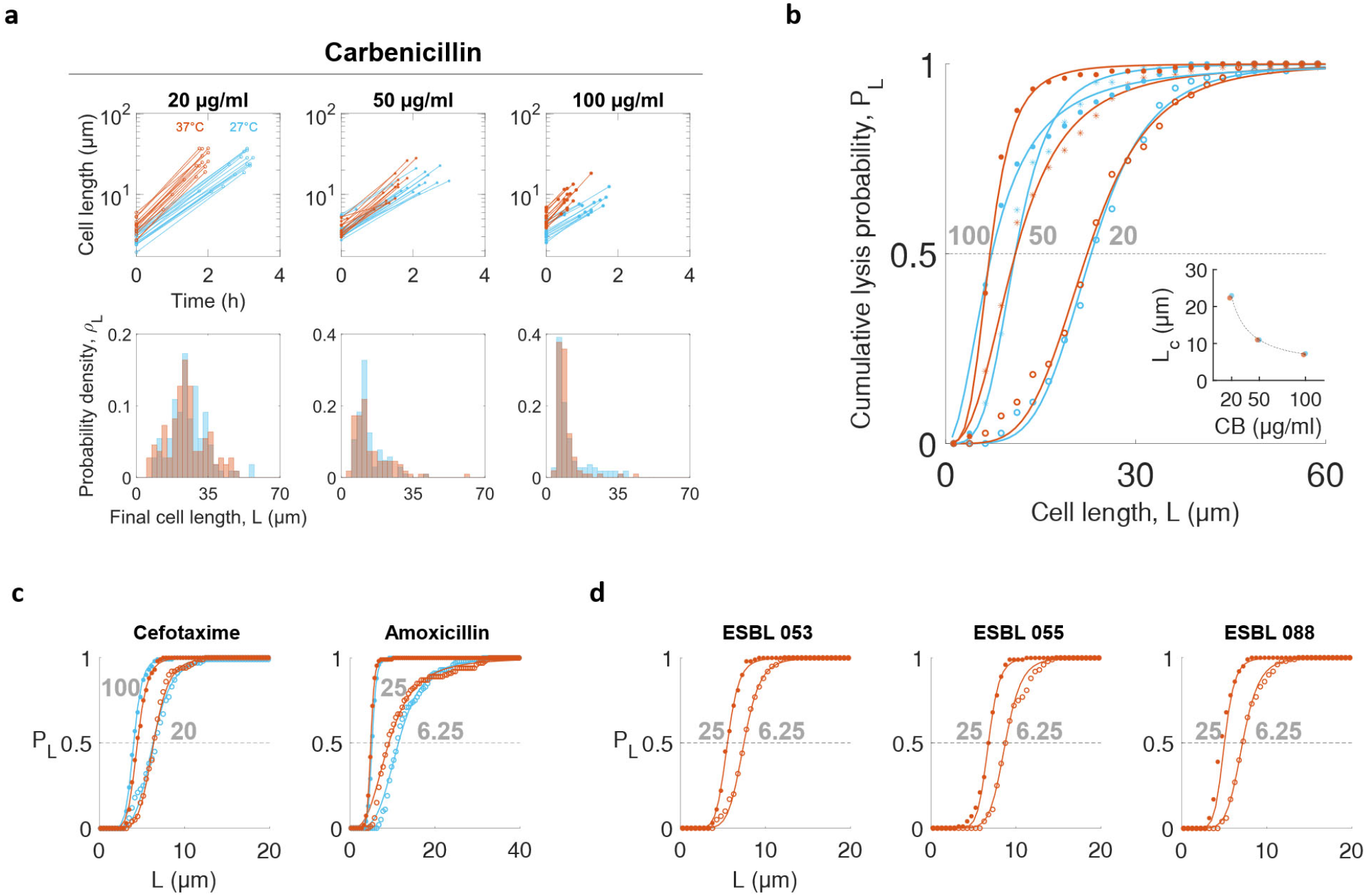
Critical lengths are shortened with an increase in antibiotic dose. **a. Probability density distributions of final lengths of MG1655 cells exposed to carbenicillin**. Individual cell lengths were measured from populations exposed to three doses of carbenicillin (20, 50, and 100 μg/ml) at two temperatures (27°C in blue and 37°C in red). Line plots show the measured initial and final lengths over time in log scale, and the first 15 cells were picked in each dataset for presentation among *n* ≥ 100 of the tracked cells in each condition (top panels, see **Table S1** for *n*). Probability density distributions (*ρ*_*L*_) of final lengths were collected with the same number of bins within the same final cell length range (bottom panels). The peak of *ρ*_*L*_ shifted to the left with increasing carbenicillin dose. **b. The probability of lysis increased with the filamentation length at each condition**. *P*_*L*_ was calculated from *ρ*_*L*_ in panel **a** (filled circle: 100 μg/ml, star: 50 μg/ml, and lined circle: 20 μg/ml). Solid lines represent fits using a log-logistic distribution (*R* > 0.99. See **Table S1** for fitted parameters). *L*_*C*_ was insensitive to temperature (27 °C in blue and 37 °C in red, carbenicillin doses in gray text), but was inversely correlated with carbenicillin (CB) dose (panel **b** inset: 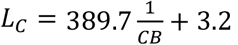, unit shown in the figure, *R*^2^ = 0.9991). **c. The probability of lysis increased with the filamentation length of bacteria treated with other beta-lactams**. MG1655 cells were treated with cefotaxime (20 and 100 μg/ml) and amoxicillin (6.25 and 25 μg/ml, shown in gray text) at two temperatures (27 °C in blue and 37 °C in red). **d. The probability of lysis increased with the filamentation length of *E. coli* pathogens treated with a beta-lactam**. *E. coli* pathogens expressing extended beta-lactam resistance were treated with amoxicillin (6.25 or 25 μg/ml, shown in gray text) and clavulanic acid (50 μg/ml) simultaneously at 37 °C. Cells were sensitized by clavulanate acid, which inhibits beta-lactamases.

We further tested if the length dependence of *P*_*L*_ and inverse correlation of *L*_*c*_ to antibiotic dose were maintained in various conditions: different beta-lactams, bacterial strains, and growth media. First, we observed the same dependence of *P*_*L*_ on cell length when using two other beta-lactam antibiotics (cefotaxime at 20 and 100 μg/ml and amoxicillin at 6.25 and 25 μg/ml) at two temperatures (27 and 37 °C) (**Figure 2c** and **Figure S3a-b**). For each antibiotic, the inverse correlation was consistently observed. Second, we found that the same trends were maintained for three clinical isolates of pathogenic *E. coli* strains (**Figure 2d** and **Figure S3c**). These isolates are resistant to beta-lactams due to their ability to express extended-spectrum beta-lactamases (ESBL). We inhibited this resistance by using clavulanic acid, a well-established beta-lactamase (Bla) inhibitor, thus rendering the isolates susceptible to beta-lactams. Specifically, these isolates were exposed to two amoxicillin doses (6.25 and 25 μg/ml) in the presence of clavulanic acid (50 μg/ml) at 37 °C. *P*_*L*_ of the isolates were also sigmoidal with the extent of filamentation, and *L*_*c*_ was shorter at higher antibiotic dose. Finally, we observed that the inverse correlation was maintained in different growth media (**Figure S4**). The final length decreased with increasing carbenicillin dose regardless of growth medium, though the absolute final lengths differed across media.

As a plausible interpretation of the dependence of *P*_*L*_ on *L*, we hypothesized that cell wall integrity remains until the total damage caused by the antibiotic accumulates (in association with elongation) to a threshold to trigger lysis (**Figure S5a**, see **Supplementary information** for details of the model). The model formulation is based on the current understanding of the beta-lactam-induced cross-linking failure which causes cell wall bursting at varying lengths ^17, 23, 27, 28^. This model draws inspiration from cooperative binding in gene transcription activation, along with a physical model of cell wall crack formation from the accumulation of peptide defects ^32^. Briefly, this model generates a gamma distribution of final length with a given threshold number (*α*) and a damage rate (*β*), which stands for the damage capacity per unit length and may increase with the increase in antibiotic concentration. The sigmoidal cumulative distribution of the gamma distribution may explain the experimentally observed sigmoidal *P*_*L*_. The parameters of each distribution–*α* and *β* of gamma distribution and *L*_*c*_ and *H* of Hill equation–are correlated: *H* is well approximated by 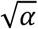 and *L*_*c*_ by 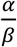 (see **Supplementary information** for detailed derivations). Therefore, *L*_*c*_ and *H* may provide simple proxies for the mechanical aspects of single-cell lysis.

### Predicting temporal dynamics of population-level responses

To map single-cell and population-level responses, we simulated and tracked the elongation and lysis of single cells in a population using a simple stochastic model (**Figure S6**, see **Methods** for model details). We simulated cell elongation with a constant (but cell-specific) exponential growth rate (*μ*) ^21^ and calculated *P*_*L*_ (using Eq. 1) and 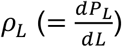 at each cell length. For each cell that has not lysed, the instantaneous lysis rate at length *L* is called the hazard function ^33, 34^, which is determined by *P*_*L*_ and *ρ*_*L*_:

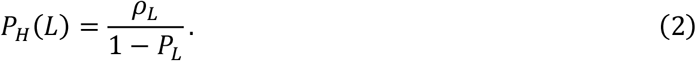

We present the formal definition of the hazard function and the derivation of Eq. 2 in the **Supplemental Information**. Briefly, Eq. 2 captures the following relationship: the probability of observed lyses between lengths *L* and *L* + *dL* is equal to the probability of cells having reached length *L* multiplied by the probability for these cells to lyse between *L* and *L* + *dL*. That is, *ρ*_*L*_*dL* = (1 − *P*_*L*_)*P*_*H*_(*L*)*dL. P*_*H*_(*L*) also allows us to compute how a population size changes as the function of the average cell length during antibiotic treatment (see below).

In this model, we assumed cell division fully stops in the presence of antibiotics but elongation continues ^12, 31^. Therefore, our stochastic single-cell length simulations show that the total biomass of the population exhibits a characteristic transient increase before a decline, while the cell number monotonically decreases over time (**Figure S6b**). These results have been observed in *in vitro* assays ^16, 24^. We define the characteristic time point when the total biomass decreases to that of the initial point as the effective treatment duration (*τ*_*E*_), as it represents the duration of antibiotic treatment sufficient to suppress the target population, despite the continued elongation of surviving cells.

Using a coarse-grained deterministic model, population biomass dynamics can be approximated by accounting for the average cell length as a function of *t, L*(*t*), and the total cell number as a function of *L, N*(*L*):

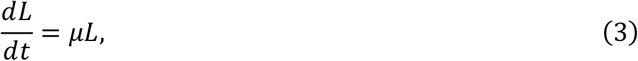

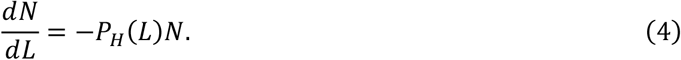

Note that the derivative in Eq. 4 is with respect to length, to follow the definition of *P*_*H*_(*L*). From Eq. 3 and Eq. 4, we can derive the temporal dynamics of total surviving biomass, which should be proportional to *LN. LN* thus follows (see **Methods** for derivation):

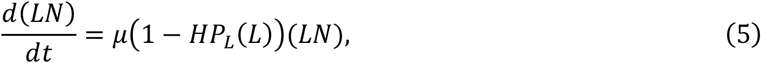

which is defined by *H, P*_*L*_, and *μ*. We note that *P*_*L*_, over time, increases and converges to 1 due to the definition of cumulative probability and the further elongation of cells.

This deterministic model allows us to compute how the total biomass changes as a function of time for different antibiotic doses (**Figure 3a**). The temporal biomass dynamics of the coarse-grained model (**Figure S7**) were quantitatively consistent with those generated from the single-cell simulations mentioned above (**Figure S6**), which started with heterogeneous initial sizes and elongation rates. Our coarse-grained model analytically demonstrates the time- and dose-dependent population biomass dynamics (**Figure 3b**). At the boundary condition, where total biomass is equal to the initial biomass (*N*(*τ*_*E*_)*L*(*τ*_*E*_) = *N*(0)*L*(0)), *τ*_*E*_ increases with an increasing *L*_*c*_ or a decreasing *μ*. Precisely, *τ*_*E*_ satisfies the following:

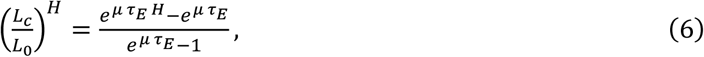

**Figure 3.**
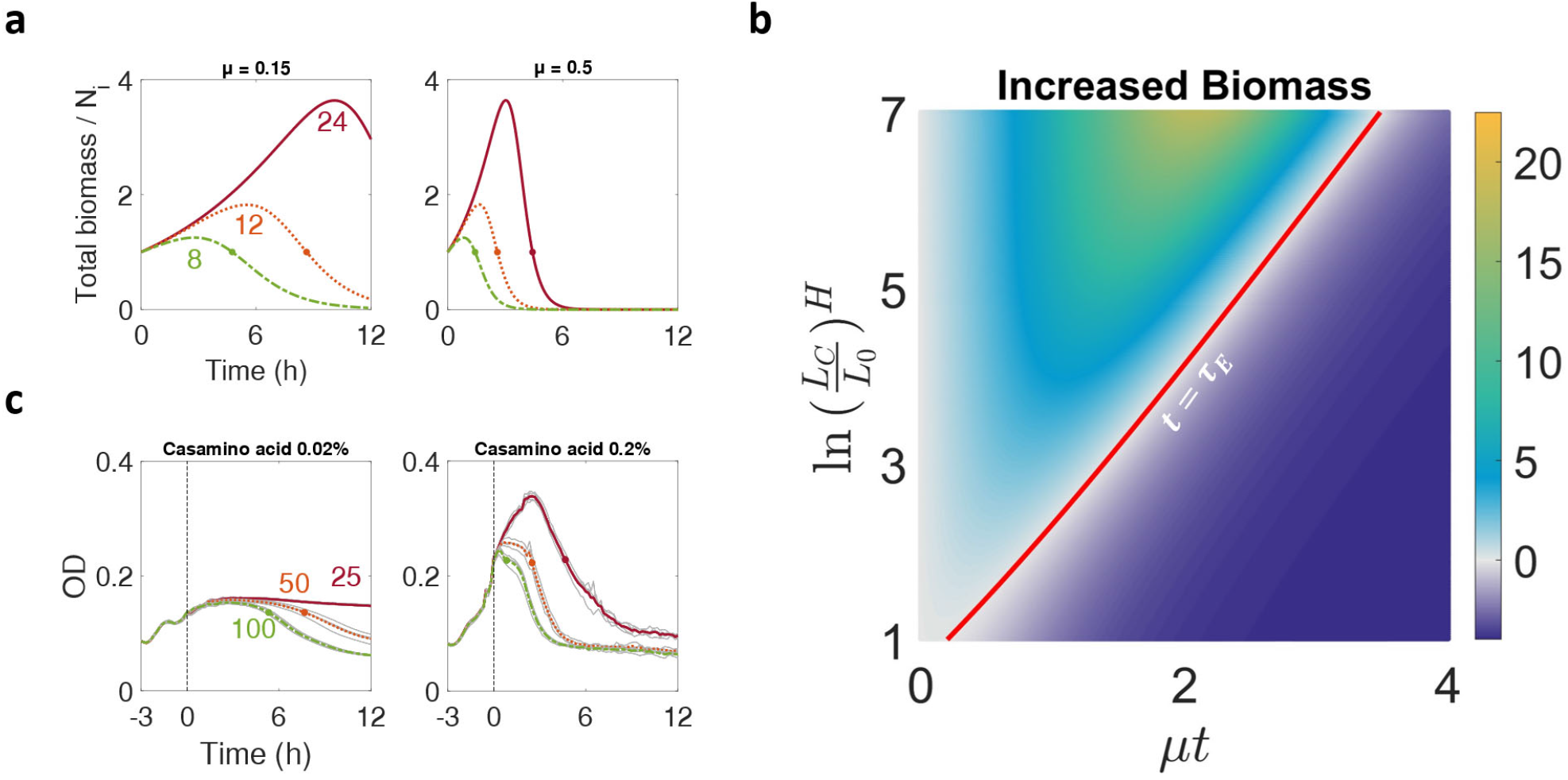
Single-cell lysis profiles predict temporal dynamics of population growth and lysis. **a. Sample simulations of the temporal dynamics of total biomass**. Three population simulations of different *L*_*C*_ (marked near each plot, arbitrary unit) with slow (left panel) and fast (right panel) elongation rates are shown in time. The effective elongation duration point (*τ*_*E*_), the point at which the total biomass is equal to the initial biomass, is depicted by a dot. The *τ*_*E*_ occurs earlier in the population either with the faster elongation rate or shorter critical length. **b. The boundary of effective elongation condition depends on *L***_***C***_ **and *τ***_***E***_. Increased biomass is shown as the color of the heatmap. The boundary condition of *τ*_*E*_ with respond to *μ* and *L*_*C*_ is plotted in red, where the increased biomass is zero after initial points. **c. Experimentally observed *τ***_***E***_ **points qualitatively verify the boundary condition of *τ***_***E***_. For growth rate modulation, casamino acid at 0.02% (left panel) and 0.2% (right panel) were added to minimal media. Cells were first cultured without carbenicillin for 3h (dashed line) and then exposed to carbenicillin (antibiotic doses are marked near each plot, μg/ml). The average of four technical repeats (gray lines) was plotted in a colored line. *τ*_*E*_ points were marked by finding the first time point at which the averaged OD became equal to or less than the OD at the start of carbenicillin exposure.

where *L*_0_ is the initial cell length. Intuitively, 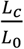 sets the critical factor to which cells can elongate before lysis while *L*_*c*_ decreases with increasing antibiotic dose. *e*^*μt*^ indicates the extent of elongation. If *t* > *τ*_*E*_, the time duration of antibiotic treatment has exceeded the limit set by 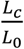. The larger *μ* is, the smaller *τ*_*E*_ would be for the same 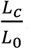, and thus the sooner the antibiotic will suppress the total biomass. Indeed, the immediate cell division and regrowth from elongated cells after few hours of antibiotic exposure has been reported ^12^, which supports that the antibiotic did not thoroughly suppress the population in a short time. More importantly, filament length was roughly proportional to the number of daughter cells, following the conservation of biomass between filamented cells and their daughters ^31^.

We experimentally assessed the dependence of *τ*_*E*_ on *L*_*c*_ and *μ*, using different concentrations of carbenicillin and casamino acids to modulate *L*_*c*_ and *μ*, respectively (**Figure 3c**). Indeed, longer *τ*_*E*_ was observed in lower antibiotic concentrations and in more slowly elongating populations. Our results establish a quantitative correlation between antibiotic dose and minimum killing time of the population.

### Predicting the linear correlation between the population-level growth and lysis rates

Eq. 5 recapitulates the two salient features of population dynamics exposed to beta-lactams mentioned above. First, according to Eq. 5, the total population biomass will increase until (1 − *HP*_*L*_) < 0, which explains time-delayed lysis. As long as *H* > 1, 1 − *HP*_*L*_ will eventually become negative, as *P*_*L*_ approaches 1. Second, the term *μHP*_*L*_ in Eq. 5 is the effective lysis rate of the population; it approaches *μH* as *P*_*L*_ approaches 1. Thus, Eq. 5 predicts a proportionality between the maximum growth rate (*G* = *μ*) and the maximum lysis rate (*D* = *μH*), with *H* being the coefficient. This proportionality is exact if there is no cell-cell variability in initial cell lengths and growth rate of each cell. However, even when such variability is considered, our numerical simulations (**Figure 4a, b**) indicate that the proportionality is maintained.

**Figure 4.**
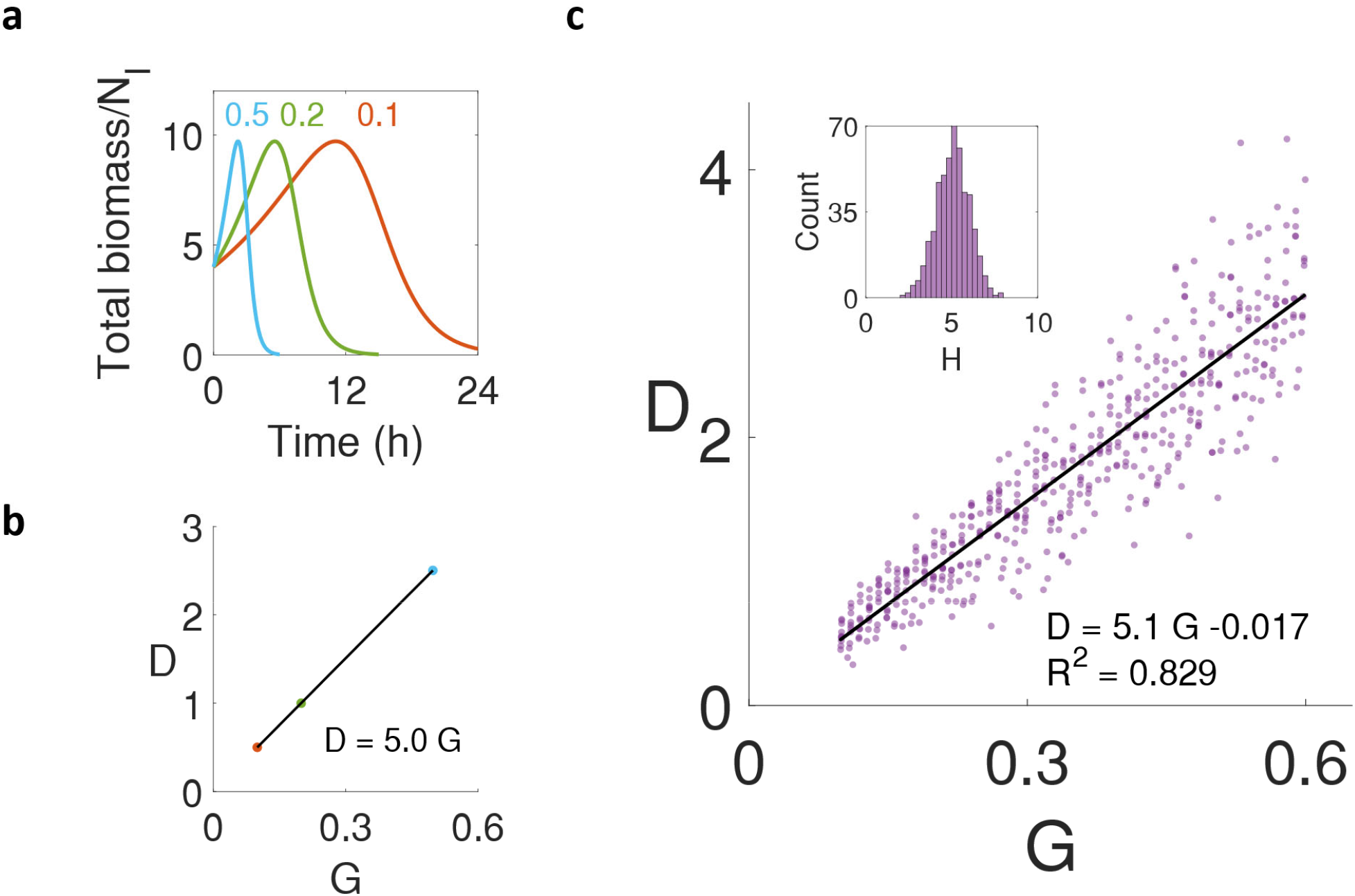
Linear correlation of maximum growth and lysis rate of the population. **a. Population biomass simulations in modulating elongation rates**. Elongation rate modulation simulations were done with a controlled initial cell number (10,000 cells), initial length (4), and *P*_*L*_ (*L*_*c*_ = 16, *H* = 5). Temporal dynamics of normalized total biomass with constant elongation rates (0.1, 0.2, and 0.5) show a decrease in population biomass at different lysis rates. **b. The maximum lysis rate increases by *H* for each maximum growth rate**. Linear regression of the maximum growth rate (G) and lysis rate (D) of each simulation in panel **a** shows the slope as 5.0, which perfectly matched the input *H*. **c. Lysis rate is linearly correlated to the growth rate**. The linearity stays with the normalized distribution of *H* (inset histogram in panel **c** with a mean of 5), which was randomly introduced to the population simulations. The slope (5.1) of the linear regression was similar to the mean of randomized *H*.

A caveat of these simulations and of Eq. 5 is the assumption of *H* being constant at different growth rates. Our experimental measurements (**Figure 2** and **Table S1**) suggest moderately variable *H*. To test the effect of this variability, we conducted numerical simulations by using normally distributed *H* values (with a mean of 5 and a variance of 1). We then collected maximum growth (*G*) and lysis rates (*D*) of each population and conducted linear regression. Despite the variability in *H*, the proportionality between lysis rate and growth rate was approximately maintained; the slope (5.1) of the linear correlation was close to the mean *H* value (**Figure 4c**).

Therefore, our model provides a simple, single-cell-based explanation for the emergence of a linear correlation between maximum population growth rate and lysis rate from previous studies ^5, 6^. We noticed that the fitted *H* values of the *P*_*L*_ in **Figure 2** were larger than those from previous experiments ^6^. On the one hand, this discrepancy reflects a potential limitation of our simplified model in quantitively matching the experimental data. On the other, it could also reflect a limitation in the resolution of the experimental data. For example, the residual biomass of lysed cells contributes to the optical density measurements, which could lead to an underestimation of the maximum lysis rates. Altogether, our results suggest that *H* poses a theoretical cap of lysis rates for different growth rates despite the variability. These results are along with our damage accumulation model, where the *H* approximates the threshold damage number (*α*) that may originate from biological mechanisms. A bigger *α* leads to a sharper response with a higher *H* (**Figure S5**).

### Predicting the non-monotonic dependence of survivor cell lengths on antibiotic doses

Our measurements show that *L*_*c*_ decreases with increasing antibiotic dose (**Figure 2b, inset**), which is likely due to the need to accumulate sufficient defects in the cell wall before it collapses. However, if the antibiotic dose is sub-lethal and the lysis probability therefore remains low, antibiotic exposure results in cells elongating longer on average before division or lysis relative to untreated cells. As such, for increasing antibiotic doses, we predicted that the average length of survivors would first increase and then decrease, that is, survivor length is biphasic (**Figure 5a**). Previous studies ^21, 35-38^ have reported this non-monotonic length dependence in antibiotic susceptibility testing but do not offer a mechanistic explanation.

**Figure 5.**
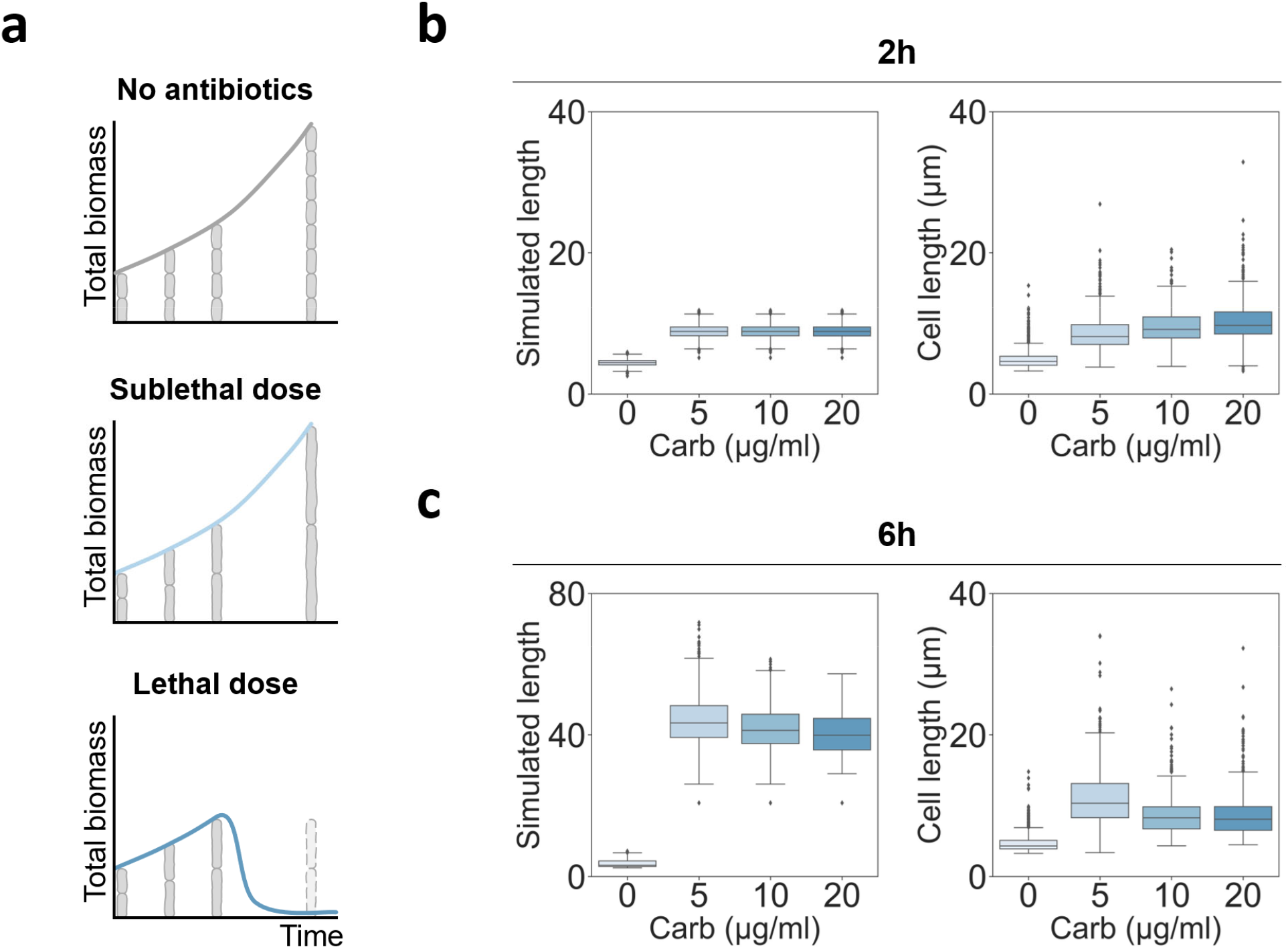
Single-cell lysis kinetics predict dose-response survivor lengths at a sublethal dose. **a. Illustration of single cells and total biomass of populations during antibiotic treatment**. At the population level, total biomass measurements of the sublethal dose antibiotic-treated population often match those of a non-treated population. Single-cell measurements, however, can distinguish the presence of antibiotics due to the elongation of individuals. Therefore, time-point single-cell measurements offer more detailed information about the lethality of antibiotics than population-level measurements. **b-c. Stochastic simulations and measurements of survivor length**. Simulated survivor lengths were plotted using the predicted *L*_*C*_ of lower doses of carbenicillin (left panels) at 2h (**b**) and 6h (**c**). The corresponding single-cell length measurements *in vitro* were plotted (right panels). In the simulation, only non-antibiotic treated cells were set to divide, when the cell length reaches the division length in the linear model (*L*_*d*_ = *aL*_*i*_ + *b*, a = 0.871, b = 2.7. Parameters were extracted from the original paper^39^). In both simulations and measurements, cells in carbenicillin-treated conditions were further elongated than control cells at all time points. At 2h exposure, antibiotic-exposed cells showed similar lengths across antibiotic concentrations since lysis is not highly probable for short filaments. At 6h exposure, survivor lengths decreased with increasing carbenicillin dose.

To examine this hypothesis, we extrapolated *L*_*c*_ of sublethal doses from the inverse correlation in **Figure 2b** and simulated length distributions of populations using our stochastic model mentioned above. We also ran separate cell length simulations for untreated cells, where cell division was enabled, using a previously established model ^39^ (**Figure 5b-c**). After a short exposure (2h), simulated cell lengths were similar across antibiotic doses because of low lysis probability for those short filaments (**Figure 5b**, left). After a prolonged exposure (6h), the length of the survivors (cells with an intact cell wall) decreased with increasing drug concentration (**Figure 5c**, left). This is mainly because cell lysis is more probable at higher doses due to the higher cumulative lysis probability at a given length. Therefore, our simulation results recapitulated the biphasic cell length trend when increasing antibiotic concentration in the sublethal range and may explain the possible kinetics that causes the trend.

To experimentally verify the model predictions, we treated cells in liquid culture with low doses of carbenicillin (0, 5, 10 μg/ml for sublethal and 20 μg/ml for lethal, **Figure S8**) and imaged every 2 hours starting from the time point when the antibiotic was added (**Figure S9**). Indeed, our single-cell simulations captured the biphasic trend of survivor lengths only in longer antibiotic treatment: after 2h, the average lengths of surviving cells were similar for different antibiotic doses, and they were longer than in the absence of the antibiotic (**Figure 5b**, right); after 6h, the biphasic dependence of average length of surviving cells on the antibiotic dose emerged (**Figure 5c**, right). We note that the quantitative aspects of experimental data, the absolute length of survivors, may differ from the simulated results due to extrapolation or different incubation conditions (liquid vs solid agarose gel).

## Discussion

Antibiotic-induced bacterial filamentation and population dynamics have not been quantitatively correlated despite the emphasis on the role of morphological protection in bacterial survival against environmental stresses and host immune systems ^40, 41^. Our single-cell measurements reveal a robust dependence of lysis probability on cell length during antibiotic treatment. In particular, the cumulative observed lysis probability over length (*P*_*L*_) showed a sigmoidal curve. We have shown that two coarse-grained empirical parameters, *L*_*c*_ and *H*, characterize the *P*_*L*_, which was unique to cell strains, growth conditions, and the type and dose of antibiotics. These two parameters serve as a quantitative basis for interpreting well-documented population-level responses to beta-lactams.

*L*_*c*_ represents the critical length for single-cell filamentation. At a given antibiotic dose, *L*_*c*_ does not change with elongation rate. It thus provides a simple explanation for why faster-growing cells are more susceptible to antibiotics: they reach *L*_*c*_ earlier. Moreover, *L*_*c*_ serves as the single-cell basis for the time-delayed killing by beta-lactams at the population level. *L*_*c*_ and *μ* define the effective treatment duration (*τ*_*E*_), which demonstrates the minimal duration needed for an antibiotic dose to effectively suppress bacterial growth (**Figure 3**). From the bacterial perspective, however, it suggests a potential phenotypic benefit of filamentation during antibiotic treatment; therefore, *τ*_*E*_ also represents the effective elongation duration for the bacterial population to survive the antibiotic treatment. Indeed, previous studies have shown that an antibiotic-treated population can recover upon the removal of the antibiotic ^10, 29, 30^. Since filamented cells produce daughter cells in proportion to their length ^31^, the continued filamentation can allow populations to recover biomass more quickly if an antibiotic is removed before *τ*_*E*_. That is, the phenotypic limit of single-cell filamentation dictates the beneficial time window for the population.

*L*_*c*_ also represents the lethality of the drug or susceptibility of bacteria in a given condition. Since *L* was inversely correlated to antibiotic dose, our results suggest that 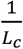 can directly report the killing capacity in a quantitative manner (**Figure 2**). The specificity of *L*_*c*_ allows comparisons of antibiotic efficacy across different antibiotics and bacterial strains. For example, if all conditions except the antibiotic agent remain identical, the agent with a shorter *L*_*c*_ is more detrimental to bacteria; if all conditions except bacterial strain remain the same, the strain with a shorter *L*_*c*_ is more vulnerable to the given antibiotic treatment condition.

The Hill coefficient (*H*) of *P*_*L*_ reflects the cell-to-cell variations in antibiotic killing kinetics. It is remarkable how the population-level metric (the linear correlation between growth and lysis rates) ^6^ quantitatively connects with a single-cell metric, *H* (**Figure 5**). The Hill coefficient measures the steepness of the response curve, and in *P*_*L*_, *H* shows how tightly the lysis is determined by cell length. Therefore, *H* can report the amount of cell-to-cell length variations in antibiotic-induced lysis. Since antibiotic resistance is often associated with the phenotypic heterogeneity-driven variance of single-cell lysis dynamics ^42, 43^, *H* can be a useful metric in projecting the resistance emergence in a population.

Additionally, our damage accumulation model suggests that *H* can emerge from underlying molecular events: the threshold number of defects (α) in the cell wall that triggers the collapse of the cell wall (**Figure S5a**). Our analysis showed that *L*_*c*_ is inversely correlated to damage capacity per length (β), which increases with the antibiotic dose. That is, our work reveals the fundamental constraint of possible molecular mechanisms that underlie single-cell responses, as illustrated by our proposed damage accumulation model.

Together, our work provides the single-cell lysis kinetics and the analytical relationship between the kinetics and population biomass dynamics using coarse-grained modeling. This relationship supports the quantitative understanding of antibiotic killing kinetics and effective treatment design. According to our further analysis, the single-cell lysis probability measures mechanical properties of the potential cell lysis mechanism (damage accumulation) during antibiotic treatment. This provides a new perspective on the fundamental understanding of cell lysis mechanism.

## Methods

### Bacterial strain

We used an *E. coli* MG1655 strain constitutively expressing a fluorescence protein from a plasmid (p15A-pTet-sfGFP-linker-Tdimer-kanR). The fluorescence was used for image analysis. We integrated the sfGFP-linker-Tdimer cassette (sfGFP-linker-Tdimer) ^44^ into the vector plasmid (p15A-pTet, kanR) by Gibson assembly. Additionally, we used isolates that were identified as ESBL-producing *E. coli* in a library from Duke Hospital’s Division of Infectious Diseases (DICON 053, 055, and 088) ^45^.

### Growth media, chemicals, and OD measurement

Unless otherwise noted, we picked a single colony from an LB plate and cultured it in 3 ml of M9CA media with 0.4% glucose overnight (∼16-18 hours). For imaging, we diluted the overnight culture in fresh media (1:10), incubated it for 2h, and used 1 μl of the diluted culture for time-lapse microscopy. All cultures were incubated in test tubes and placed in a 37 °C shaker with 225 rpm.

For time-course population-level measurements, we washed and resuspended overnight cultures in PBS and diluted to make 0.1 of OD_600_ in fresh M9 media containing either 0.2% or 0.02% casamino acids at the same rate, using 96-well plates (Corning) with a sterile transparent seal (ThermalSeal RTS, Excel Scientific) to prevent evaporation. All population dynamics were measured using Tecan Infinite Pro200, where OD_600_ was measured every 10 minutes with 10-second orbital shaking before each measurement. All single-cell experiments and most population measurements were done at 27 °C or 37 °C. Some population-level dynamics were also measured at 30 °C (**Figure 3c** and **Figure S8**), which was used to further tune overall population growth rates.

When applicable, kanamycin at 50 μg/ml (Sigma) to select for plasmid-containing cells and carbenicillin (Genesee Scientific), amoxicillin (Sigma), cefotaxime (Sigma), and/or clavulanic acid (Sigma) with appropriate concentrations were added to the growth media.

### Time-lapse microscopy

We prepared new 1.5% agarose growth media gel ahead of every imaging experiment. In the 15ml conical tube, UltraPure Low Melting Point Agarose (Invitrogen) powder was dissolved into 3 ml growth media by putting it in a 70°C water bath for 3 min and mixed well by pipetting. We aliquoted 1 ml of the solution in microtubes, added antibiotics, gently mixed, and loaded into small wells made with adhesive isolators [either Geneframe (Thermo Fisher, 25μl) or Press-To-Seal Silicon Isolator (Grace Bio-Labs, Ø 8mm × 0.8mm depth)] on a clean glass slide. The solution was flattened with another glass slide and solidified at room temperature for 5 minutes. Pre-culture (1 μl) was loaded onto the gel, spread by tilting, allowed to sit for 3 min for cell setting, and covered with a coverslip.

A Keyence microscope (BZ-X710 and BZ-X800) with an incubation chamber for a microscope (INU Tokai hit) was set up for time-lapse imaging. We used a 40X objective for phase-contrast images and when applicable, including a DsRed filter for fluorescent images. Images were taken at 5-minute intervals, and 7 levels of focus at 0.7 μm z-stack intervals were investigated to determine focus with the focus tracking option of the software. Most focused images were automatically chosen by the microscope software (BZ-X analyzer) for image analysis. The incubation chamber was set up with temperature control only: to achieve x°C, the top was set to (x+10)°C and the bottom to (x+2)°C according to the manufacturer’s instruction. We used either 27°C or 37°C as x for time-lapse microscopy.

### Image analysis of final length from time-lapse microscopy and lysis probability fitting

Manually drawn line segment length was measured using Fiji ^46^ to capture the long-axis length of elongated and curved cells. Length measurements were mainly done with phase-contrast images, with additional red fluorescence channel images being used only when cell outlines were faint in phase-contrast images. We used MATLAB codes to convert units and generate probability distributions. The MATLAB R2021a curve fitting tool was used for ill equation fitting of cumulative probability distributions.

### Mathematical model of mapping single-cell lysis to population biomass dynamics

From Eq. 2-4, we can derive the *temporal* dynamics of total cell number:

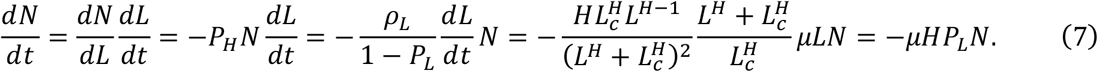

Further derivation of total biomass then follows:

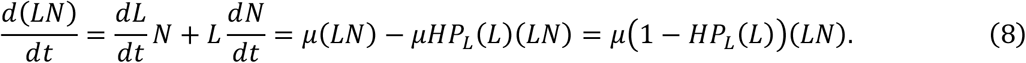

Assuming an initial cell length of *L*_*0*_ and an initial cell number of *N*_*0*_, *L*(*t*) and *N*(*t*)can be derived from Eq. 1-3 and 7 by following:

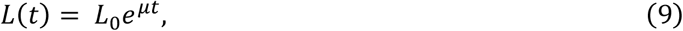

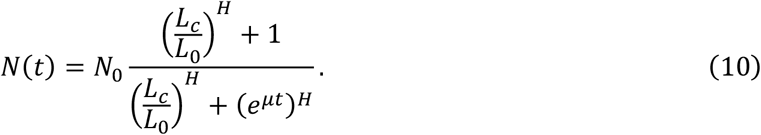

### Mathematical model simulations

We used MATLAB R2021a for numeric simulations. The codes associated with Figs. 3, 5, and S6 are provided.

In stochastic simulations, we initialized and ran simulations of the model as follows:

1. For a population, set *L*_*c*_ and *H* of *P*_*L*_ and initialize 2,000 cells (*N*_*i*_) with an average initial length (*L*_*i*_) and an average single-cell elongation rate (*μ*).
2. Add Gaussian noise to *L*_*i*_ and μ of each cell, but not to exceed 10% of the given average values, and ensure *L*_*i*_ > 0 and *μ* > 0.
3. For cells without division, compute *L*(*t*) = *L*_*i*_*e*^*μt*^, *ρ*_*L*_ *L*(*t*)), and *P*_*L*_ *L*(*t*)). To integrate the hazard funtion lysis rate (*P*_*H*_) within simulation time step size, we used *P*_*H*_ *L*(*t*)) *L*(*t*) − *L*(*t* − Δ*t*)) as hazard function. By generating one random number (*d*), uniform between 0 and 1, for each cell, we set *L*(*t* ≥ *T*) = 0 when *P*_*H*_ *L*(*t*)) > *d*, which implies lysis with no residual biomass.
4. For cells with division under non-treated conditions, force a cell to divide into two cells of equal length after reaching the designated division length (*L*_*d*_ = *aL*_*i*_ + *b*) according to the linear model ^39^. We tracked up to a maximum of 10,000 cells.
5. Compute total biomass by summing up *L*(*t*) of all cells.

In deterministic simulations, we used an averaged initial length 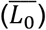 and elongation rate 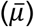. Eq. 3 and 7, to yield *L*(*t*) and *N*(*t*), were computed with ode45. Total biomass was then the product of those two as shown in Eq. 5. In the exponential growth and lysis rate simulations, the net growth rate of a population was calculated as the log of total biomass divided by the time interval, 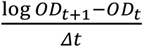. In each simulation, the maximum growth rate was found from the maximum net growth rate, and the maximum lysis rate was found by subtracting the minimum from the maximum net growth rate.

### Survivor length measurments

Overnight cultures were split into new culture tubes. We added carbenicillin to the culture in proper doses and incubated the tubes in the 37 °C shakers with 225 rpm. We loaded 1 μl of the culture onto a glass slide and covered. Due to low cell number, cells treated with 20 μg/mL carbenicillin at the 6-hour timepoint were spun down (100 μL, 2000 rcf, 2 min) and resuspended in 20 μL of the same media for imaging. 40X phase contrast images from 3 locations of each experimental condition were taken to yield more than 100 cells to be analyzed. Cell lengths were extracted with customized single-cell segmentation Python code based on the scikit-image package ^47^.

## Author contributions

KK conceived the research, performed both modeling and experimental analysis, and wrote the manuscript. TW, ES, BL, and VA assisted in mathematical modeling and data interpretation. HM assisted with experimental analysis and manuscript revisions. LY conceived the research, assisted in research design and modeling, and wrote the manuscript. All authors approved the manuscript.

## Acknowledgment

We thank Caroline Connor for assistance in editing the manuscript and Allison J. Lopatkin for valuable advice in finalizing the manuscript. This work was partially supported by the National Institutes of Health (L.Y., R01AI125604, R01GM110494, and R01EB029466), and the National Science Foundation (L.Y., MCB-1937259). The funders had no role in study design, data collection and analysis, decision to publish, or preparation of the manuscript.

## Notes

### Competing Interest Statement

The authors have declared no competing interest.

## Reference

1. Hamad, B. The antibiotics market. Nat Rev Drug Discov 9, 675–676 (2010).

2. Wong, F. & Amir, A. Mechanics and Dynamics of Bacterial Cell Lysis. Biophys J 116, 2378–2389 (2019).

3. Craig, W.A. Interrelationship between pharmacokinetics and pharmacodynamics in determining dosage regimens for broad-spectrum cephalosporins. Diagn Microbiol Infect Dis 22, 89–96 (1995).

4. Craig, W.A. Pharmacokinetic/Pharmacodynamic Parameters: Rationale for Antibacterial Dosing of Mice and Men. Clinical Infectious Diseases 26, 1–12 (1998).

5. Tuomanen, E., Cozens, R., Tosch, W., Zak, O. & Tomasz, A. The rate of killing of Escherichia coli by beta-lactam antibiotics is strictly proportional to the rate of bacterial growth. Journal of general microbiology 132, 1297–1304 (1986).

6. Lee, A.J. et al. Robust, linear correlations between growth rates and beta-lactam-mediated lysis rates. Proceedings of the National Academy of Sciences of the United States of America 115, 4069–4074 (2018).

7. Meredith, H.R., Lopatkin, A.J., Anderson, D.J. & You, L. Bacterial Temporal Dynamics Enable Optimal Design of Antibiotic Treatment. PLOS Computational Biology 11, e1004201 (2015).

8. Buijs, J., Dofferhoff, A.S.M., Mouton, J.W., Wagenvoort, J.H.T. & van der Meer, J.W.M. Concentration-dependency of β-lactam-induced filament formation in Gram-negative bacteria. Clinical Microbiology and Infection 14, 344–349 (2008).

9. Elliott, T.S.J. & Greenwood, D. The response of Pseudomonas aeruginosa to azlocillin, ticarcillin and cefsulodin. Journal of Medical Microbiology 16, 351–362 (1983).

10. Chen, K., Sun, G.W., Chua, K.L. & Gan, Y.-H. Modified Virulence of Antibiotic-Induced <em>Burkholderia pseudomallei</em> Filaments. Antimicrobial Agents and Chemotherapy 49, 1002–1009 (2005).

11. Mason, D.J., Power, E.G., Talsania, H., Phillips, I. & Gant, V.A. Antibacterial action of ciprofloxacin. Antimicrob Agents Chemother 39, 2752–2758 (1995).

12. Cayron, J., Dedieu, A. & Lesterlin, C. Bacterial filament division dynamics allows rapid post-stress cell proliferation. bioRxiv, 2020.2003.2016.993345 (2020).

13. Kjeldsen, T.S.B., Sommer, M.O.A. & Olsen, J.E. Extended spectrum β-lactamase-producing Escherichia coli forms filaments as an initial response to cefotaxime treatment. BMC microbiology 15, 63–63 (2015).

14. Paulander, W. et al. Bactericidal antibiotics increase hydroxyphenyl fluorescein signal by altering cell morphology. PloS one 9, e92231–e92231 (2014).

15. Cushnie, T.P.T., O’Driscoll, N.H. & Lamb, A.J. Morphological and ultrastructural changes in bacterial cells as an indicator of antibacterial mechanism of action. Cellular and Molecular Life Sciences 73, 4471–4492 (2016).

16. Eng, R.H., Cherubin, C., Smith, S.M. & Buccini, F. Inoculum effect of beta-lactam antibiotics on Enterobacteriaceae. Antimicrobial agents and chemotherapy 28, 601–606 (1985).

17. Cho, H. et al. Bacterial cell wall biogenesis is mediated by SEDS and PBP polymerase families functioning semi-autonomously. Nature Microbiology 1, 16172 (2016).

18. Chung, H.S. et al. Rapid beta-lactam-induced lysis requires successful assembly of the cell division machinery. Proceedings of the National Academy of Sciences of the United States of America 106, 21872–21877 (2009).

19. Vigouroux, A. et al. Class-A penicillin binding proteins do not contribute to cell shape but repair cell-wall defects. eLife 9, e51998 (2020).

20. Choi, J. et al. A rapid antimicrobial susceptibility test based on single-cell morphological analysis. Science Translational Medicine 6, 267ra174–267ra174 (2014).

21. Rolinson, G.N. Effect of β-Lactam Antibiotics on Bacterial Cell Growth Rate. Microbiology 120, 317–323 (1980).

22. Spratt, B.G. Distinct penicillin binding proteins involved in the division, elongation, and shape of Escherichia coli K12. Proceedings of the National Academy of Sciences of the United States of America 72, 2999–3003 (1975).

23. Yao, Z., Kahne, D. & Kishony, R. Distinct single-cell morphological dynamics under beta-lactam antibiotics. Mol Cell 48, 705–712 (2012).

24. Zahir, T. et al. Image-Based Dynamic Phenotyping Reveals Genetic Determinants of Filamentation-Mediated β-Lactam Tolerance. Frontiers in Microbiology 11 (2020).

25. Cho, H., Uehara, T. & Bernhardt, Thomas G. Beta-Lactam Antibiotics Induce a Lethal Malfunctioning of the Bacterial Cell Wall Synthesis Machinery. Cell 159, 1300–1311 (2014).

26. Burdett, I.D. & Murray, R.G. Septum formation in Escherichia coli: characterization of septal structure and the effects of antibiotics on cell division. Journal of bacteriology 119, 303–324 (1974).

27. Daly, K.E., Huang, K.C., Wingreen, N.S. & Mukhopadhyay, R. Mechanics of membrane bulging during cell-wall disruption in Gram-negative bacteria. Physical Review E 83, 041922 (2011).

28. Şimşek, E. & Kim, M. Power-law tail in lag time distribution underlies bacterial persistence. Proceedings of the National Academy of Sciences 116, 17635–17640 (2019).

29. El Meouche, I., Siu, Y. & Dunlop, M.J. Stochastic expression of a multiple antibiotic resistance activator confers transient resistance in single cells. Scientific Reports 6, 19538 (2016).

30. Zahir, T. et al. High-throughput time-resolved morphology screening in bacteria reveals phenotypic responses to antibiotics. Communications Biology 2, 269 (2019).

31. Wehrens, M. et al. Size Laws and Division Ring Dynamics in Filamentous Escherichia coli cells. Current biology : CB 28, 972-979.e975 (2018).

32. Huang, K.C., Mukhopadhyay, R., Wen, B., Gitai, Z. & Wingreen, N.S. Cell shape and cell-wall organization in Gram-negative bacteria. Proceedings of the National Academy of Sciences 105, 19282–19287 (2008).

33. Gompertz, B.P. XXIV. On the nature of the function expressive of the law of human mortality, and on a new mode of determining the value of life contingencies. In a letter to Francis Baily, Esq. F. R. S. &c. Philosophical Transactions of the Royal Society of London, 513–583.

34. Collett, D. Modelling Survival Data in Medical Research. (Springer US, 1994).

35. Chung, C.-C. et al. Screening of Antibiotic Susceptibility to β-Lactam-Induced Elongation of Gram-Negative Bacteria Based on Dielectrophoresis. Analytical Chemistry 84, 3347–3354 (2012).

36. Oh, J. et al. Three-dimensional label-free observation of individual bacteria upon antibiotic treatment using optical diffraction tomography. Biomed. Opt. Express 11, 1257–1267 (2020).

37. McLaughlin, H.P. & Sue, D. Rapid antimicrobial susceptibility testing and β-lactam-induced cell morphology changes of Gram-negative biological threat pathogens by optical screening. BMC microbiology 18, 218–218 (2018).

38. Fredborg, M. et al. Automated image analysis for quantification of filamentous bacteria. BMC microbiology 15, 255–255 (2015).

39. Tanouchi, Y. et al. A noisy linear map underlies oscillations in cell size and gene expression in bacteria. Nature 523, 357–360 (2015).

40. Khan, F., Jeong, G.-J., Tabassum, N., Mishra, A. & Kim, Y.-M. Filamentous morphology of bacterial pathogens: regulatory factors and control strategies. Applied Microbiology and Biotechnology (2022).

41. Justice, S.S., Hunstad, D.A., Cegelski, L. & Hultgren, S.J. Morphological plasticity as a bacterial survival strategy. Nature Reviews Microbiology 6, 162–168 (2008).

42. Andrews, J.M. Determination of minimum inhibitory concentrations. J Antimicrob Chemother 48 Suppl 1, 5–16 (2001).

43. Artemova, T., Gerardin, Y., Dudley, C., Vega, N.M. & Gore, J. Isolated cell behavior drives the evolution of antibiotic resistance. Molecular systems biology 11, 822–822 (2015).

44. Xia, A. et al. Dual-Color Fluorescent Timer Enables Detection of Growth-Arrested Pathogenic Bacterium. ACS Infectious Diseases 4, 1666–1670 (2018).

45. Kanamori, H. et al. Genomic Analysis of Multidrug-Resistant Escherichia coli from North Carolina Community Hospitals: Ongoing Circulation of CTX-M-Producing ST131-<i>H</i>30Rx and ST131-<i>H</i>30R1 Strains. Antimicrobial Agents and Chemotherapy 61, e00912–00917 (2017).

46. Schindelin, J. et al. Fiji: an open-source platform for biological-image analysis. Nature Methods 9, 676–682 (2012).

47. van der Walt, S. et al. scikit-image: image processing in Python. PeerJ 2, e453 (2014).

