## Supplementary Information for "Mapping single-cell responses to population-level dynamics during antibiotic treatment"

| Strain | Antibiotics | Dose (μg/ml) | Temp. (°C) | nCells | $L_c$ (μm) | H | R-square |
|---|---|---|---|---|---|---|---|
| MG1655 | Carbenicillin | 20 | 27 | 110 | 22.97 | 5.00 | 0.9980 |
| | | | 37 | 110 | 22.33 | 4.45 | 0.9948 |
| | | 50 | 27 | 104 | 10.96 | 3.96 | 0.9992 |
| | | | 37 | 110 | 10.99 | 2.69 | 0.9961 |
| | | 100 | 27 | 110 | 7.25 | 2.07 | 0.9914 |
| | | | 37 | 106 | 6.97 | 3.72 | 0.9984 |
| | Cefotaxime | 20 | 27 | 100 | 6.30 | 5.76 | 0.9993 |
| | | | 37 | 100 | 6.37 | 7.08 | 0.9998 |
| | | 100 | 27 | 100 | 4.03 | 8.15 | 0.9999 |
| | | | 37 | 100 | 4.47 | 7.55 | 0.9999 |
| | Amoxicillin | 6.25 | 27 | 100 | 11.36 | 4.08 | 0.9977 |
| | | | 37 | 100 | 9.12 | 2.97 | 0.9946 |
| | | 25 | 27 | 100 | 5.16 | 10.44 | 1.0000 |
| | | | 37 | 110 | 4.85 | 11.38 | 1.0000 |
| ESBL053 | Amoxicillin | 6.25 | 37 | 100 | 7.46 | 7.80 | 0.9997 |
| | | 25 | | 100 | 5.51 | 8.04 | 0.9999 |
| ESBL055 | | 6.25 | | 100 | 8.74 | 9.23 | 0.9997 |
| | | 25 | | 100 | 6.75 | 10.48 | 0.9999 |
| ESBL058 | | 6.25 | | 100 | 7.16 | 7.36 | 0.9998 |
| | | 25 | | 100 | 5.04 | 8.39 | 0.9999 |

**Table S1. Fitted results of $P_L$ in antibiotic dose and temperature modulation assays.**

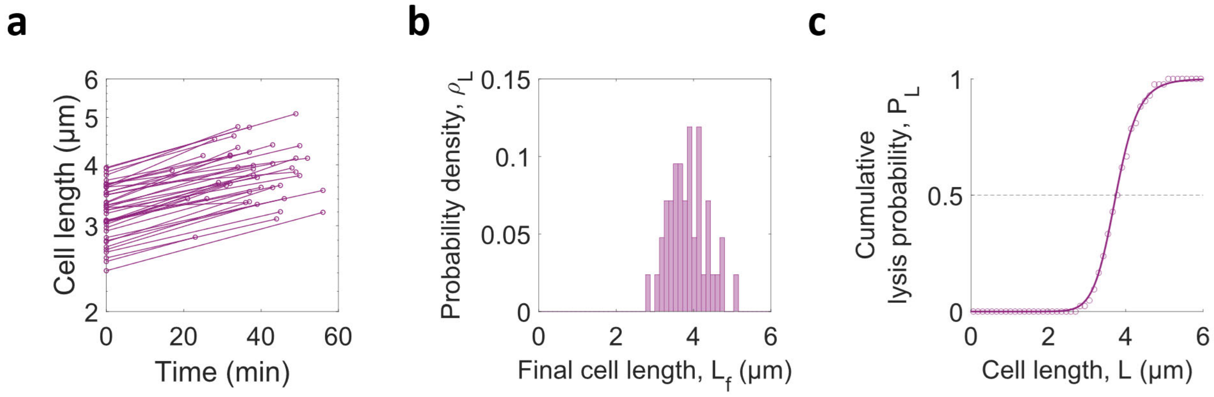


**Figure S1. D-cycloserine-induced lysis probability in a single cell increased with the extent of filamentation.**

**a. Initial and final lengths of cells over time.** From the raw data, only cells that elongated and lysed after treatment were chosen for analysis. Line plots show log scaled initial and final length over time.

**b. Probability density distribution from final length.** Probability density distribution ($\rho_L$) over the final length was taken by normalizing probability distributions.

**c. Cumulative lysis probability increases with elongation during treatment**. Cumulative probabilities ($P_L$) of the $\rho_L$ are shown in dot plot. Solid line shows the log-logistic fitted $P_L$.

**a**

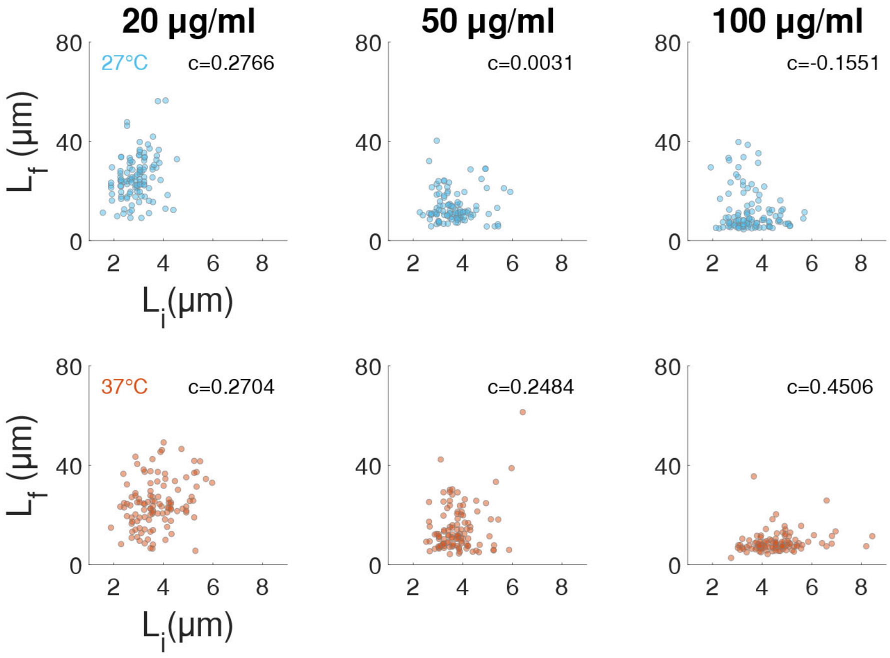


**b**

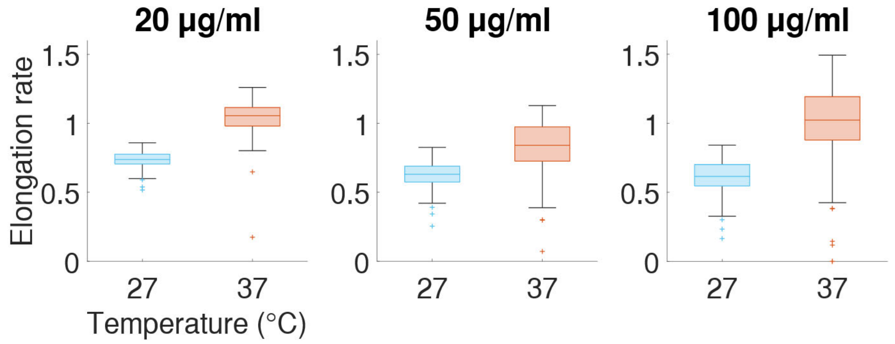


**Figure S2. Effects of initial cell length and incubation temperature on carbenicillin-induced elongation.**

**a. Final length is independent from initial length.** Scatter plots of final length over initial length show weak correlations between initial and final lengths in all conditions. $c$ is the correlation coefficient as a measure of linear independence and printed on the top right of each panel.

**b. Temperature modulates elongation rate.** Individual elongation rates were calculated with initial and final length measurements of individual cells. In accordance with the slopes in **Figure 2a**, elongation rate was higher at 37 °C vs. 27 °C.

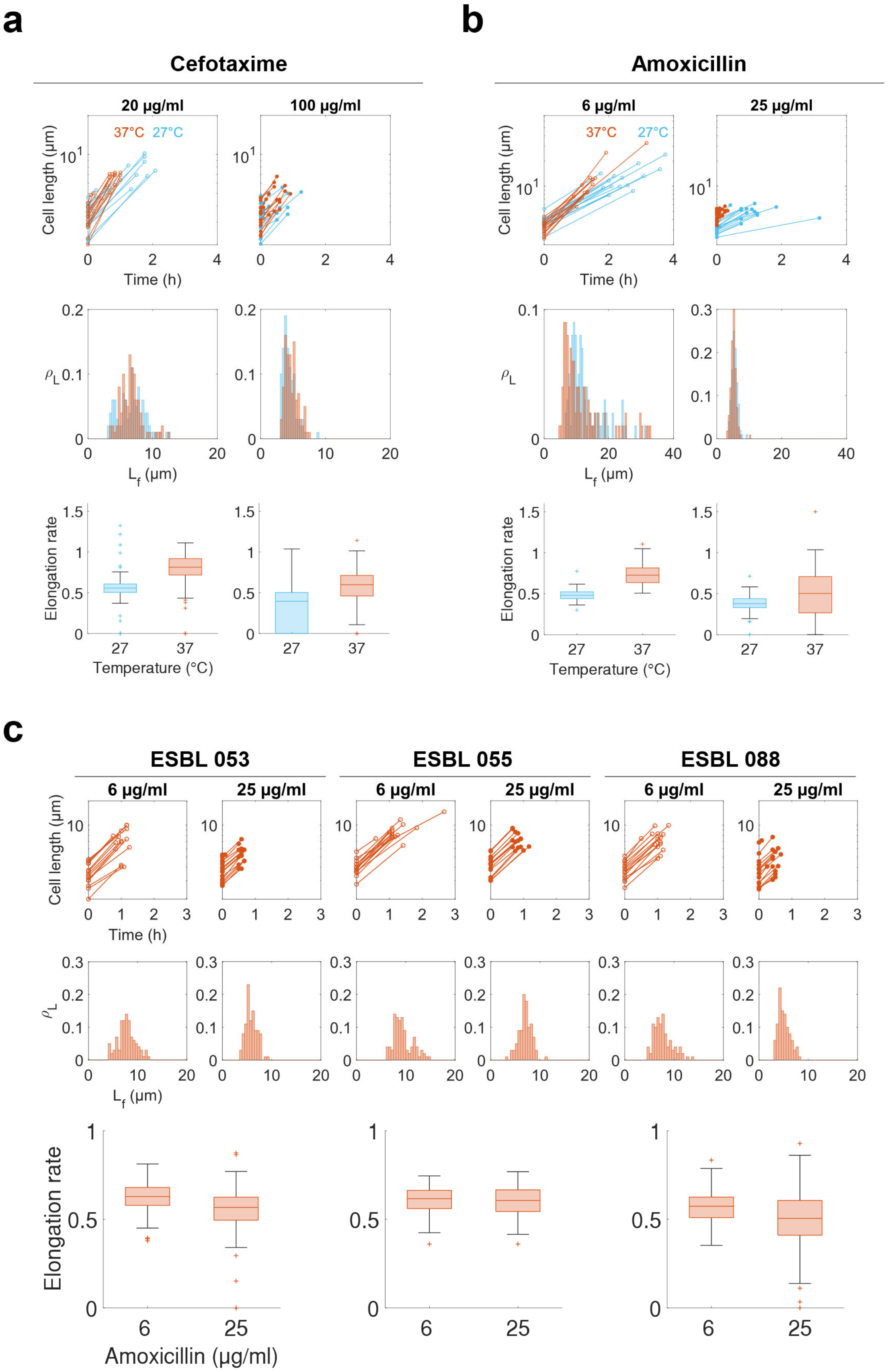
a
b
Cefotaxime
Amoxicillin
20 µg/ml
100 µg/ml
6 µg/ml
25 µg/ml
37°C
27°C
Cell length (µm)
Time (h)
L_f (µm)
ρ_L
Elongation rate
Temperature (°C)
c
ESBL 053
ESBL 055
ESBL 088
Amoxicillin (µg/ml)

**Figure S3. Final length measurements for finding $P_L$ in other beta-lactams and ESBL-producing strains.**

**a-b. Cefotaxime and amoxicillin treatment in *E. coli* MG1655 with temperature modulation.** Two doses of cefotaxime (20 and 100 μg/ml) and amoxicillin (6.25 and 25 μg/ml) at two temperatures (27°C in blue and 37°C in red) were tested. Initial and final lengths of the first 15 cells are shown in the first row for display. Lysis probability densities from the final lengths of all measured cells are shown in the second row. Elongation rates of all measured cells are shown in the third row.

**c. Amoxicillin-treated ESBL-producing *E. coli* from the patient isolate library with Bla inhibition.** Clavulanate acid (50 μg/ml) was added with amoxicillin (6.25 and 25 μg/ml) to the clinical isolates and incubated at 37 °C. Cell length, probability density, and elongation rate plots are derived by the same methods in panels **a** and **b**. At the later stage of low-dose amoxicillin treatment, these cells formed localized swelling [1, 2]—a gradually inflated body from the middle of its cylindrical region—and the lab strain formed a bulb through the cell wall without changing the cell width. Therefore, we note that the final length of amoxicillin treated (6.25 μg/ml) cells could be slightly longer than the measured length, because the biomass that contributed to extending the width may have extended length if cells did not form local swell.

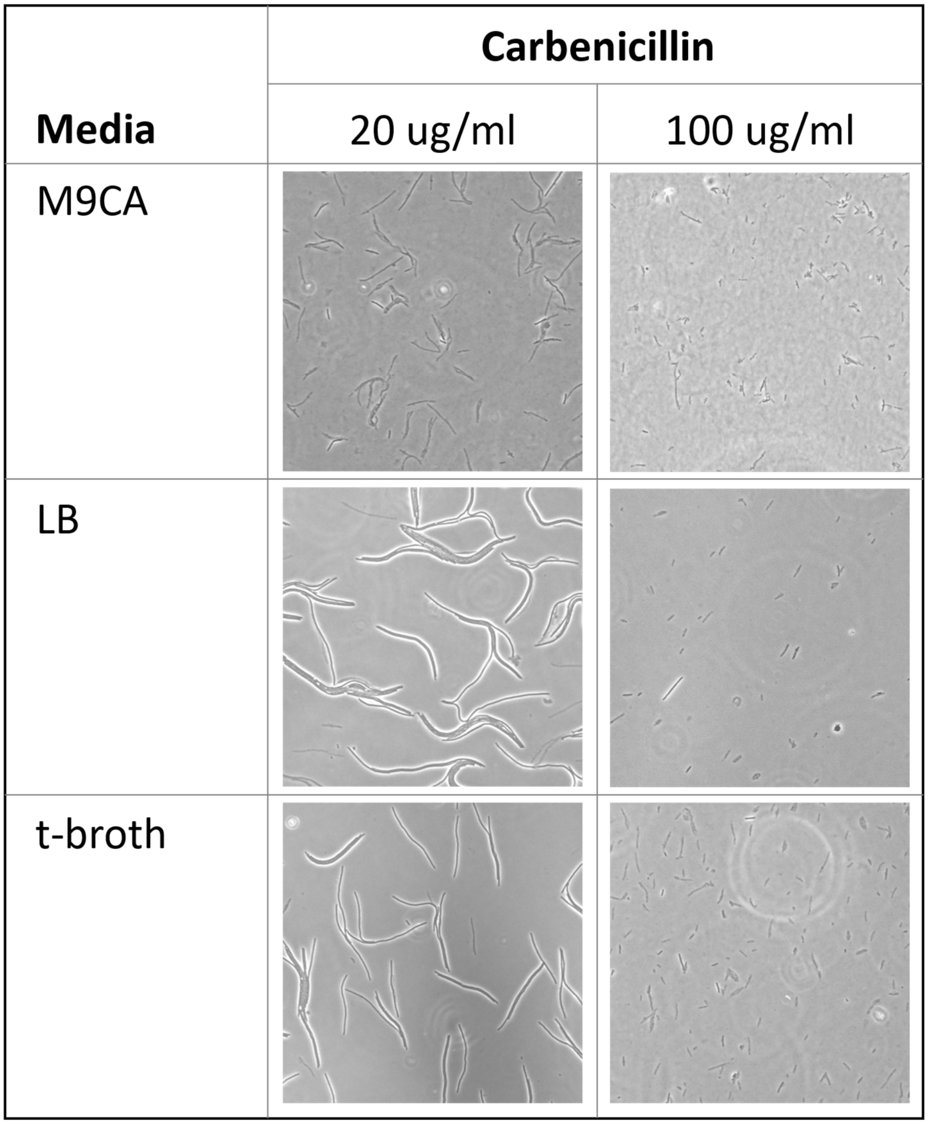


**Figure S4. Dose-dependency of $L_c$ maintained in different growth media.**

Exponentially growing cells were loaded on different media gel with the same absolute carbenicillin doses for time-lapse microscopy. Images taken after around 6 to 8 hours of antibiotic exposure were picked for displaying low-dose carbenicillin-treated cells. The inverse relationship of $L_c$ and carbenicillin dose was conserved in each medium. Compared to cells grown in M9CA medium, used in **Figure 2** experiments, the lengths of the cells grown in LB and t-broth were longer; therefore, $L_c$ is sensitive to growth media.

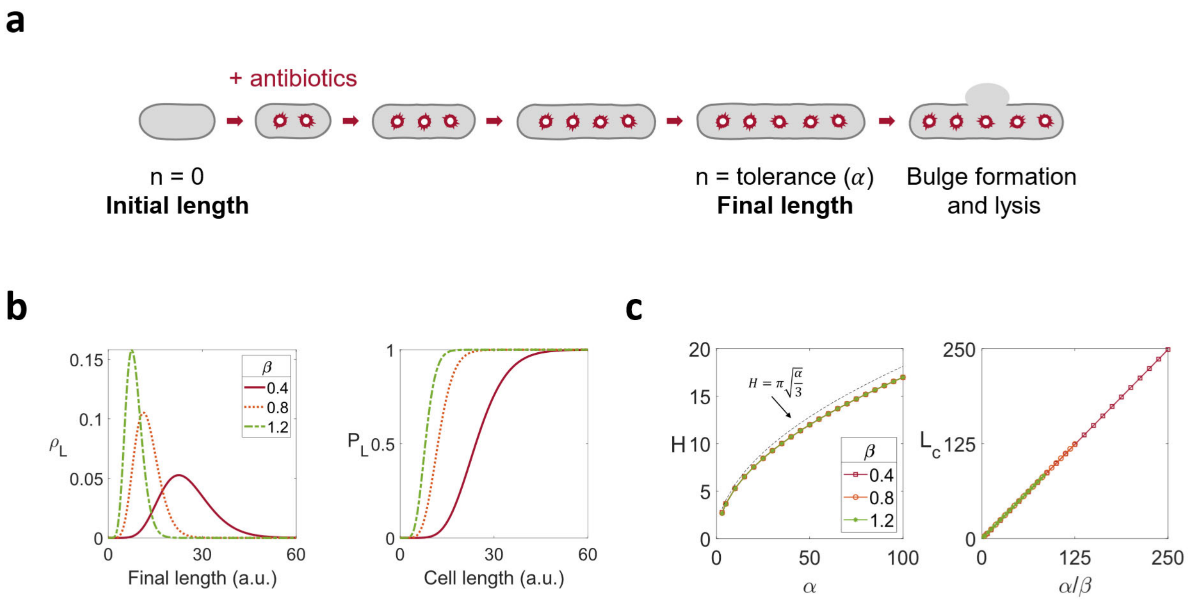


**Figure S5. Damage accumulation model provides a plausible interpretation of the log-logistic distribution through gamma distribution.**

**a. Damage accumulates on the cell wall until lysed.** A schematic diagram of the damage accumulation model for cell lysis. Upon addition of antibiotics, a cell is assumed to accumulate damage on its cell wall during elongation, with an antibiotic dose-dependent rate of β. When the total number of damages to a cell reaches a threshold value α, the cell formed a bulge and lysed in a short time.

**b. Final length follows a gamma distribution.** Under the damage accumulation model, the final length of a cell that has α damages follows a gamma distribution with the parameters of α and β (see **Supplementary Note 1**). Probability density (PDF, $\rho_L$, left) and cumulative density (CDF, $P_L$, right) of the three different rates (β = 0.4, 0.8, and 1.2) recapitulate the experimental distributions shown in **Figure 2b**. All three plots used a constant threshold (α = 10).

**c. Parameters of gamma and log-logistic distributions are correlated.** Gamma distributions that were generated with different parameter sets of α and β were fitted to log-logistic distributions ($P_L = \frac{L^H}{L^H + L_c^H}$). $H$ was not sensitive to β (left) but highly sensitive to α (bottom left). Black dashed line shows the approximation of $H$ to $\alpha$ in 2nd order. $L_C$ was proportional to α/β (right). $R^2$ of all log-logistic fits were larger than 0.997.

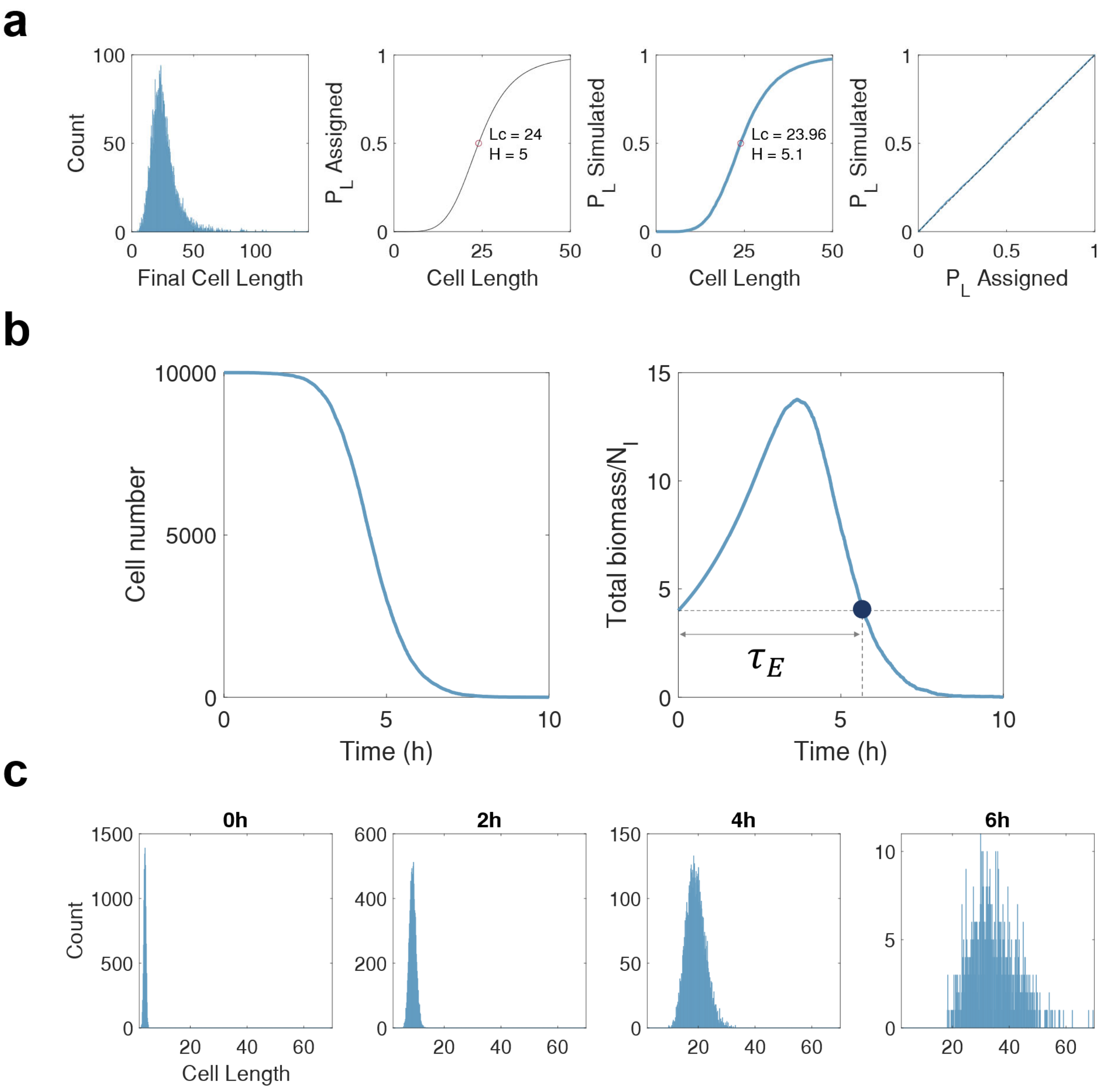


**Figure S6. Stochastic simulation confirms the single-cell level lysis profiles.**

**a. Stochastic final length simulations.** Stochastic simulation of final length distribution (the first panel) of a population with 10,000 cells was done with an assigned cumulative probability $P_A = \frac{L^H}{L^H+Lc^H}$ ($L_c$ =24 and $H$ =5, the second panel). The cumulative distribution function from the distribution, simulated $P_L$ (the third panel), was achieved and compared to the assigned $P_L$ (the last panel).

**b. Simulated population responses via collection of individual responses.** Simulation results show that the cell number only decreases because the cells cannot divide but are lysed (left panel). However, the sum of the biomass of survivors (total biomass) increases due to elongation and low lysis probability during early exposure and then decreases due to higher lysis probability (right panel). The time point at which total biomass crosses the initial biomass is the effective elongation duration, $\tau_E$.

**c. Time-course cell length distributions.** Survivor length distributions at 0 to 6h elongation.

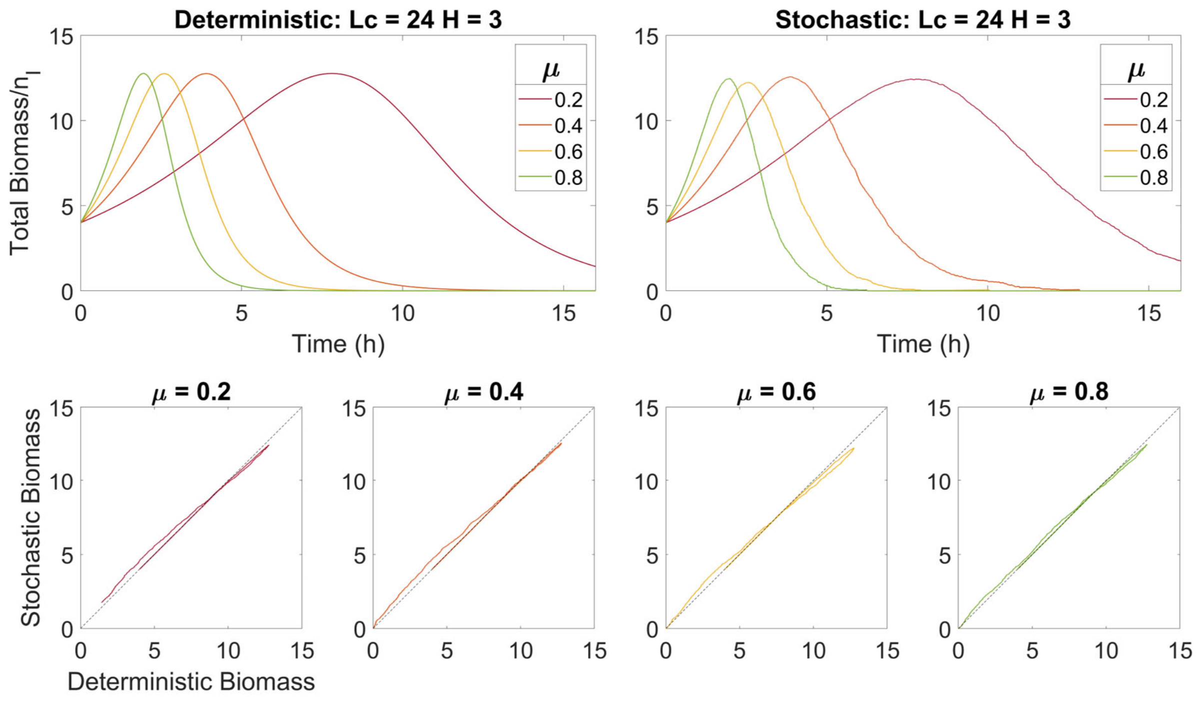


**Figure S7. Deterministic models can represent populational biomass dynamics using average parameters of stochastic models.**

Total biomass over time is shown in panel **a**, using the same average initial length and $P_L$ parameters ($L_c$ and $\mathrm{H}$) with four different average elongation rates. Deterministic and stochastic simulations were compared 1:1 by using total biomass at every time step in panel **b**.

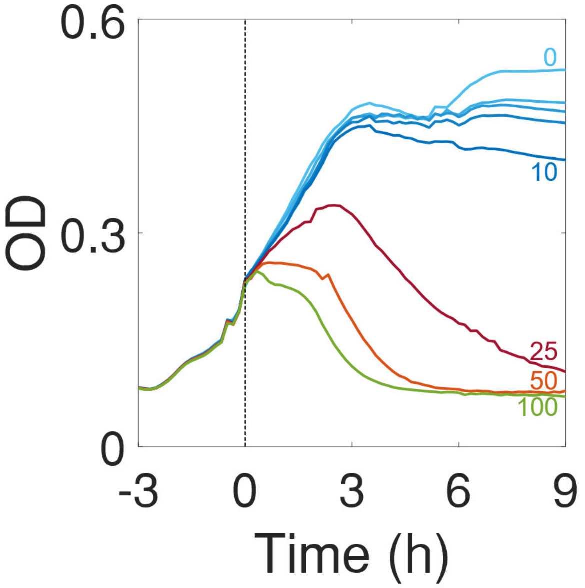


**Figure S8. Dose-response population growth and lysis dynamics.**

Carbenicillin (0, 1, 2.5, 5, 10, 25, 50, and 100 μg/ml) was added at time 0. Populations with low-dose antibiotics (≤ 10 μg/ml) show similar OD curves.

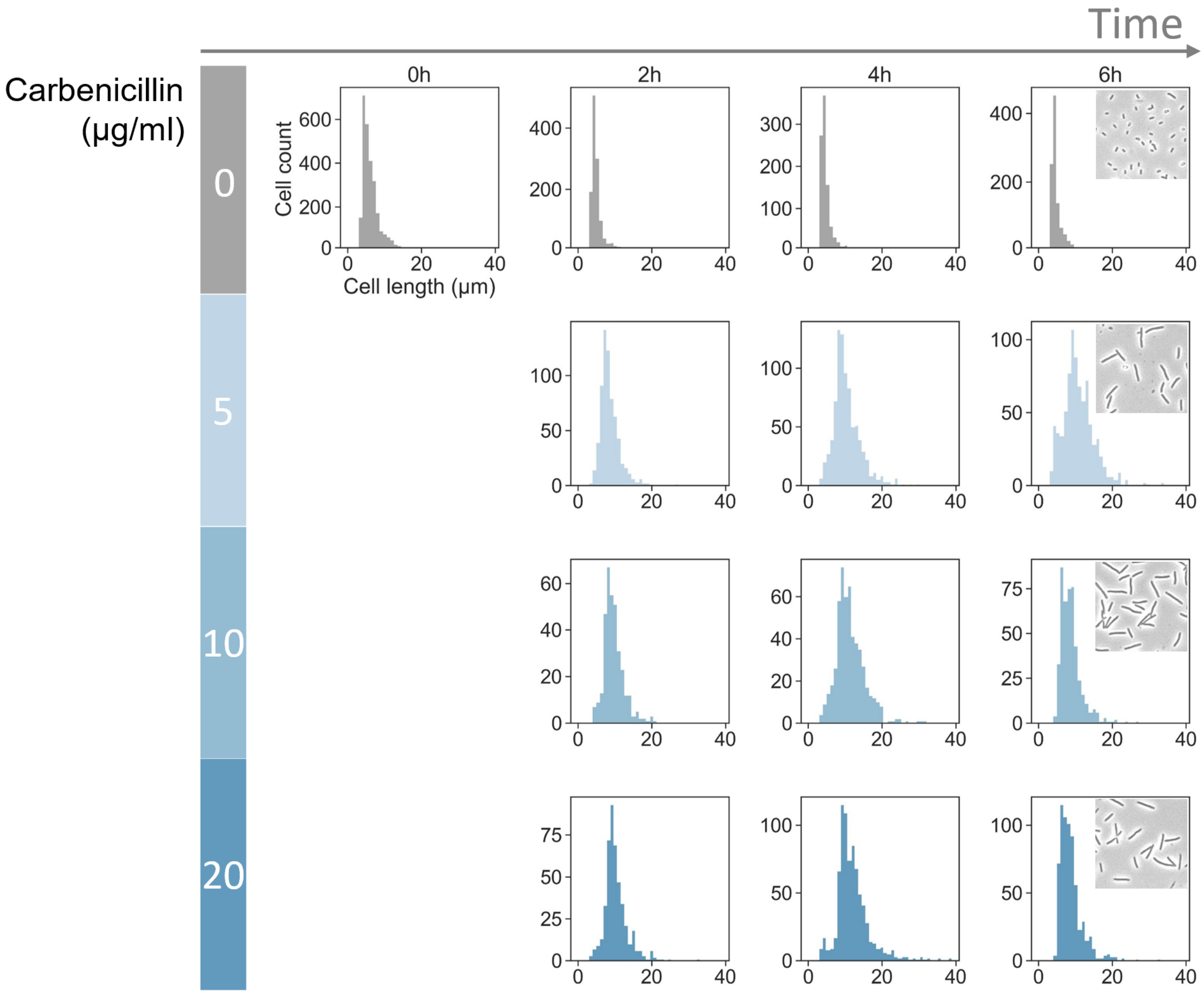


**Figure S9. Cell length distributions following carbenicillin treatment over time.**

Cell lengths were measured from the microscope images of sampled cells in time-course and dose-response experiments. The single-cell images were captured every 2 hours from the carbenicillin (0, 5, 10, and 20 μg/ml) addition at time 0. Long-axis length distributions were generated with the same number of bins.

### Model development

*Mathematical model of damage accumulation*

We assumed that 1) reaching a threshold number ($\alpha$) of unsuccessful peptidoglycan cross-linkages on the cell wall causes a loss of cell wall integrity and triggers lysis and 2) the number of failures being made per unit cell length is the damage rate, $\beta$([A]), which increases with antibiotic dose. According to our model, final length is then the length of cells when $\alpha$ failures have been acquired: its density distribution follows a gamma distribution of which the cumulative distribution is also a sigmoid.

We define the probability of having one cross-linking failure per unit length as damage rate ($\beta$) and allow only one failure in unit length. That is, for any $\Delta L \to 0$, the probability of one damage is $p_1(\Delta L) = \beta\Delta L + o(\Delta L)$, no damage is is $p_0(\Delta L) = 1 - \beta\Delta L$, and multiple damages is $p_{n\geq 2}(\Delta L) = 0$. Since we assume that the cross-linkage failure accumulates independently, an infinitesimal elongation ($dL$) of $L$-length cell without damage should follow:

$$p_0(L + dL) = p_0(L)p_0(dL) = p_0(L)(1 - \beta dL). \quad \text{(S1)}$$

Therefore,

$$\frac{dp_0(L)}{dL} = -\beta p_0(L). \quad \text{(S2)}$$

Similarly, the probability of an infinitesimal elongation ($dL$) of $L$-length cell with $n$ damages should follow:

$$p_n(L + dL) = p_{n-1}(L)p_1(dL) + p_n(L)p_0(dL) = p_{n-1}(L)\beta dL + p_n(L)(1 - \beta dL), \quad \text{(S3)}$$

$$\frac{dp_n(L)}{dL} + \beta p_n(L) = \beta p_{n-1}(L), \quad \text{(S4)}$$

$$\frac{d}{dL}\left[e^{\beta L}p_n(L)\right] = \beta e^{\beta L}p_{n-1}(L). \quad \text{(S5)}$$

where Eq. S5 is a general recursive equation for series $\{p_0(L), p_1(L), p_2(L), \cdots\}$. Combining Eq. S2 and $p_0(0) = 0$, we have $p_0(L) = e^{-\beta L}$. Then, the probability of $L$-length cell with $n$ damages is:

$$p_n(L) = \frac{(\beta L)^n}{n!} e^{-\beta L}. \tag{S6}$$

If we assume that the lysis occurs when a cell attains $\alpha$ damages, the final length distribution of cells with $\alpha$ damages ($P_L(\alpha)$) should follow:

$$P_L(\alpha) = 1 - \sum_{i=0}^{\alpha-1} p_i(L) = 1 - e^{-\beta L} \sum_{i=0}^{\alpha-1} \frac{(\beta L)^i}{i!}. \tag{S7}$$

Here, we note that Eq. S7 is the cumulative density function of final lengths and known as the Gamma distribution.

In accordance with the log-logistic fitting, the dependence of $P_L$ on $L$ in gamma distributions shifts leftward at a larger $\beta$ with constant $\alpha$ (**Figure S5a**). This trend is consistent with the experimental observations: all else being equal, a higher antibiotic dose (indicates larger $\beta$) resulted in a smaller $L_c$ (the length at the mean of cumulative distributions) (**Figure 2**).

Our numerical analysis showed that $L_c$ was proportional to $\alpha/\beta$ , and $H$ increased approximately with the square root of $\alpha$ (**Figure S5b**). These correlations emerge from the analytic solutions under the limiting case when $H \gg 1$.

Briefly, we assume the log-logistic distribution (with its CDF being the Eq 1 from direct data fitting) and the gamma distribution (arising from the damage-accumulation model) approximately describe the same data. Thus, the means and the variances of two-fitted distributions should be equal to each other:

$$\frac{L_c \frac{\pi}{H}}{sin \frac{\pi}{H}} = \frac{\alpha}{\beta}, \tag{S8}$$

$${L_c}^2 \left( \frac{\frac{2\pi}{H}}{sin \frac{2\pi}{H}} - \frac{\left(\frac{\pi}{H}\right)^2}{sin^2 \frac{\pi}{H}} \right) = \frac{\alpha}{\beta^2}. \tag{S9}$$

Further simplification of Eq. S8 shows that $L_c \approx \frac{\alpha}{\beta}$ since $\frac{\frac{\pi}{H}}{sin\frac{\pi}{H}} \approx 1$ when $H \gg 1$.

Plugging $L_c \approx \frac{\alpha}{\beta}$ into Eq. S9 gives:

$$\frac{\frac{2\pi}{H}}{sin\frac{2\pi}{H}} - \frac{\left(\frac{\pi}{H}\right)^2}{sin^2\frac{\pi}{H}} = \frac{1}{\alpha}. \tag{S10}$$

Expanding the left-hand-side terms to the second order using $\frac{x}{sinx} = 1 + \frac{1}{6}x^2 + O(x^4)$ gives:

$$1 + \frac{1}{6}\left(\frac{2\pi}{H}\right)^2 - \left(1 + \frac{1}{6}\left(\frac{2\pi}{H}\right)^2\right)^2 = \frac{1}{\alpha}. \tag{S11}$$

Further simplification of the left-hand-side terms to the second order of $H$ derives $H \approx \pi\sqrt{\frac{\alpha}{3}}$. These results indicate that measured $L_c$ and $H$ are approximately determined by the maximum number of accumulated cross-linkage failures ($\alpha$) and the damage rate ($\beta$) to retain the cell wall integrity.

*Derivation of the the hazard function formula*

The hazard function, also known as the instantaneous death rate or instantaneous failure rate, was initially introduced by Gompertz in 1825 [3]. It is often used in modeling the death of individuals in an age-distributed population, where the independent variable is time (age). However, it can be defined in terms of any monotonically increasing variable.

The Eq. 2, $P_H(L) = \frac{\rho_L}{1-P_L}$, is the hazard function formula, expressed as a function of cell length ($L$), where length is monotonically increasing variable between initial to final lengths. Below is a derivation of the hazard function formula from its definition [4].

In our study, we define $P_H(L)$ in terms of cell length ($L$):

$$P_H(L) \equiv \lim_{dL \to 0} \frac{P(L < l < L + dL | l > L)}{dL} \ . \tag{S12}$$

By definition, the hazard function describes the instantaneous lysis rate of an individual cell at cell length $L$, that has survived to $L$. The numerator of Eq. S12 is a conditional probability, which can be expanded and rearranged according to the Bayes theorem:

$$\begin{aligned} P(L < l < L + dL | l > L) &= \frac{P(l > L | L < l < L + dL) P(L < l < L + dL)}{P(l > L)} \\ &= \frac{P(L < l < L + dL)}{P(l > L)}, \end{aligned} \tag{S13}$$

as $P(l > L | L < l < L + dL)$ is always 1, reflecting certainty.

The denominator of Eq. S13 is the survival function of the population:

$$P(l > L) = 1 - cumulative\ lysis\ probability = 1 - P_L. \tag{S14}$$

Plugging Eqs. S13 and S14 into Eq. S12, we have:

$$\begin{aligned} P_H(L) &= \lim_{dL \to 0} \frac{P(L < l < L + dL)}{P(l > L)} \frac{1}{dL} \\ &= \lim_{dL \to 0} \frac{P(l < L + dL) - P(l < L)}{dL} \frac{1}{P(l > L)} \\ &= \frac{1}{P(l > L)} \lim_{dL \to 0} \frac{P(l < L + dL) - P(l < L)}{dL} \\ &= \frac{1}{1 - P_L} \frac{dP_L}{dL} \\ &= \frac{\rho_L}{1 - P_L}, \end{aligned}$$

which is the hazard function formula.

### SI Reference


1. Cushnie, T.P.T., O'Driscoll, N.H. & Lamb, A.J. Morphological and ultrastructural changes in bacterial cells as an indicator of antibacterial mechanism of action. *Cellular and Molecular Life Sciences* **73**, 4471-4492 (2016).
2. Burdett, I.D. & Murray, R.G. Septum formation in Escherichia coli: characterization of septal structure and the effects of antibiotics on cell division. *Journal of bacteriology* **119**, 303-324 (1974).
3. Gompertz, B.P. XXIV. On the nature of the function expressive of the law of human mortality, and on a new mode of determining the value of life contingencies. In a letter to Francis Baily, Esq. F. R. S. &c. *Philosophical Transactions of the Royal Society of London*, 513 - 583.
4. Collett, D. *Modelling Survival Data in Medical Research*. (Springer US, 1994).
